# A transposable element insertion in *IAA16* disrupts splicing and causes dicamba resistance in *Bassia scoparia*

**DOI:** 10.1101/2024.07.19.604363

**Authors:** Jacob S. Montgomery, Neeta Soni, Sofia Marques Hill, Sarah Morran, Eric L. Patterson, Seth A. Edwards, Sandaruwan Ratnayake, Yu-Hung Hung, Pratheek H. Pandesha, R. Keith Slotkin, Richard Napier, Franck Dayan, Todd A. Gaines

**Affiliations:** Department of Agricultural Biology, Colorado State University, Fort Collins, Colorado, USA.; Department of Plant, Soil and Microbial Sciences, Michigan State University, East Lansing, Michigan, USA; Donald Danforth Plant Science Center, St. Louis, Missouri, USA; Division of Biological Sciences, University of Missouri, Columbia, Missouri, USA; Division of Biology & Biomedical Sciences, Washington University in St. Louis, St. Louis, Missouri, USA; School of Life Sciences, University of Warwick, Coventry, UK

**Keywords:** Herbicide Resistance, Auxin, Evolution, Genetic Mapping, Transposable Elements, Weed Science

## Abstract

A dicamba-resistant population of kochia (*Bassia scoparia*) identified in Colorado, USA in 2012 was used to generate a synthetic mapping population that segregated for dicamba resistance. Linkage mapping associating dicamba injury with genotype derived from restriction-site-associated DNA sequencing identified a single locus in the kochia genome associated with resistance on chromosome 4. A mutant version of *Auxin/Indole-3-Acetic Acid 16* (*AUX/IAA16*; a gene previously implicated in dicamba resistance in kochia) was found near the middle of this locus in resistant plants. Long read sequencing of dicamba-resistant plants identified a recently inserted Ty1/Copia retrotransposon near the beginning of the second exon of *AUX/IAA16*, leading to disruption of normal splicing. A molecular marker for this insertion allows for rapid detection of resistance. Stable transgenic lines of *Arabidopsis thaliana* ectopically expressing the mutant and wildtype alleles of *AUX/IAA16* were developed. *Arabidopsis thaliana* plants expressing the mutant *AUX/IAA16* allele grew shorter roots on control media. However, transgenic root growth was less inhibited on media containing either dicamba (5 μM) or IAA (0.5 μM) when compared to non-transgenic plants or those expressing the wildtype allele of *AUX/IAA16.* In vitro assays indicate reduced binding affinity and more rapid dissociation of the mutant AUX/IAA with TIR1 in the presence of several auxins, and protein modeling suggests the substitution of the glycine residue in the degron domain of AUX/IAA16 is especially important for resistance. A fitness cost associated with the mutant allele of *AUX/IAA16* has implications for resistance evolution and management of kochia populations with this resistance mechanism.

**Significance:** Auxin mimics are amongst the most important herbicides in modern agriculture. Evolution of weeds that are resistant to these herbicides threatens sustainable crop production. Understanding the basis of auxin herbicide resistance informs the development of improved weed control technologies. Additionally, auxin-resistant mutations and their pleotropic effects help us understand auxin perception and signalling. We describe a transposable element insertion within an herbicide target site gene that alters splicing and reduces synthetic and natural auxin perception.

## Introduction

Weed control in modern agricultural production systems relies largely on herbicides. The first synthetic herbicides to be commercialized (e.g. 2,4-D; described by Quastel, 1950) were eventually determined to mimic the endogenous plant hormone auxin. The low cost of these synthetic auxin herbicides, matched with their ability to selectively control broadleaf weeds in cereal production, led to rapid adoption of herbicide-based weed management. As new herbicides were discovered and commercialized, mixing multiple herbicides in a single spray mixture has allowed for robust control of many of the most important weeds in diverse agricultural settings. While evolution of herbicide resistant weed populations has reduced the efficacy of many herbicides (Gaines et al., 2020), evolution of resistance to synthetic auxin herbicides has been slower and less frequent than for other modes of action (Busi et al., 2018; Heap, 2024). Thus, some of the oldest herbicides (e.g., 2,4-D and dicamba) remain some of the most widely used across many agricultural settings. Indeed, auxin resistance traits were engineered into dicot crops (e.g. soybean and cotton) to provide farmers more control options against weed populations that had evolved glyphosate resistance (Behrens et al., 2007; Wright et al., 2010). These traits expanded the potential use and value of two of the most used synthetic auxin herbicides, dicamba and 2,4-D. This expansion has had the unintended effect of increasing selection pressure for resistance alleles and is correlated with a recent increase in the number of resistance reports to these herbicides. Beyond weed control, these synthetic auxin compounds have been crucial for experiments to understand auxin perception and signaling, perhaps the most complex of all plant hormone signaling pathways (Ma et al., 2017).

While the physiological mechanism of action for many herbicides has been well understood for decades, the cascade of cellular responses following the application of a synthetic auxin herbicide is convoluted and is the subject of many recent studies. The topic has been reviewed several times, with new experimental results incrementally advancing our knowledge of the key factors leading to plant death (Grossmann, 2010; Song, 2014; Christoffoleti et al., 2015; McCauley et al., 2020). Auxin, whether synthetic or natural, first binds to one or more of the isoforms of its receptor protein, Transport Inhibitor Response 1/Auxin-Signaling F-Box (TIR1/AFB) (Dharmasiri et al., 2005; Kepinski & Leyser, 2005). Once bound to TIR1/AFB, auxin acts as a “molecular glue”, mediating the interaction between TIR1/AFB and the co-receptor Auxin/Indole-3-Acetic Acid (AUX/IAA) proteins. Identification of conserved domains and experimentation have produced a list of residues in TIR1/AFB and AUX/IAA that are vital for this interaction (Tan et al., 2007; Uzunova et al., 2016; Ramans-Harborough et al., 2023).

This interaction leads to the ubiquitination and rapid degradation of AUX/IAA proteins (Gray et al., 2001; Zenser et al., 2001). Because AUX/IAA proteins act as transcriptional regulators, AUX/IAA degradation leads to the modulation of auxin-responsive gene expression (Grossmann, 2010). TIR1/AFB and AUX/IAA are each multi-gene families with varying numbers of homologs. The diversity in gene members of these families allow for many possible combinations of TIR1/AFB and AUX/IAA, which in turn allow for tissue, temporal and environmental regulation of auxin responses. Different synthetic auxin herbicides likely promote the interaction of certain AUX/IAA proteins with certain TIR1/AFB proteins more efficiently, but this network of interactions is still not well understood. For example, LeClere et al. (2018) show that AUX/IAA16 preferentially binds with TIR1 and AFB6 (a homolog of AFB found in members of amaranthaceae, but not brassicaceae) in the presence of some auxin analogs, but not others.

A disruption in any step of this auxin signaling system (e.g. TIR1 binding auxin, AUX/IAA interacting with TIR1/AFB, or the ubiquitination and degradation pathway of AUX/IAA) could cause reduced activity of synthetic auxin herbicides (Ruegger et al., 1998; LeClere et al., 2018; Todd et al., 2020). Thus, evolution of synthetic auxin resistance should be relatively simple and common across weed species in the face of incredibly pervasive and strong selection pressure. However, mutants causing resistance to synthetic auxin herbicides are likely also to cause a reduction in fitness compared to the wildtype due to the conserved function of these proteins (Roux & Reboud, 2005; Wu et al., 2021), indicating that resistance mutations are likely to be purged during temporal absence of synthetic auxin herbicide selection pressure during rotations in land and herbicide use. Nevertheless, there are several reported cases of synthetic auxin herbicide resistance becoming established in some weed species, including examples of enhanced metabolic detoxification in synthetic auxin herbicide resistant weeds similar to the mechanism of selectivity in tolerant crops (Torra et al., 2024). Interestingly, in cases in which the genetic basis of resistance has been determined, all of these reports pertain to mutations in or near the degron domain of Aux/IAA genes. LeClere et al. (2018) and Ghanizadeh et al. (2024) report, in closely related species, different mutations of a specific glycine residue in AUX/IAA16 that is known to be vital to normal auxin signaling. In unrelated species, de Figueiredo et al. (2022b) report a 27-bp deletion that shortens the distance between the degron and PB1 domains of IAA2, and Krishnan et al. (2024) report a double deletion mutation flanking the degron region of IAA20.

Here, we seek to understand the genetic and molecular basis of dicamba resistance in a population of kochia (*Bassia scoparia*) collected from Colorado, USA. We use linkage mapping to identify a region of the genome that segregates with resistance and contains a mutant version of *AUX/IAA16*. Long read sequencing allowed us to identify a transposable element insertion in the coding region of *AUX/IAA16* that disrupts normal splicing and changes several amino acids in the protein product relative to the wildtype. Ectopic expression in *Arabidopsis thaliana* proves this mutant *AUX/IAA16* is sufficient to cause resistance to dicamba. Surface plasmon resonance assays detect a reduction in binding affinity and more rapid dissociation for the mutant AUX/IAA protein, and protein modeling points to the importance of a glycine-to-threonine substitution in the degron domain of this mutant. This mutation also seems to confer a fitness cost associated with this mechanism of dicamba resistance, with implications for agricultural management of dicamba-resistant weed populations.

## Results and Discussion

### Characterization of herbicide resistance

Dose response experiments were conducted to confirm and quantify the level of dicamba and 2,4-D resistance in a population of kochia (named M32) collected from Akron, Colorado, USA (Figure 1A and 1B). Parameter estimates for two-parameter log-logistic models for each population used in these experiments are shown in Table S1 and Table S2. In the dicamba experiment, populations M32 and 9425 (a known dicamba-resistant kochia population previously characterized by LeClere et al. (2018) and Pettinga et al. (2018)) had ED50 estimates 7.6 (p<0.0001) and 10.5 (p<0.0001) times greater than an herbicide-sensitive kochia population (7710; described by Preston et al. (2009)). The ED50 parameter represents the dose of herbicide needed to cause 50% injury to that specific population. These estimates were significantly different from the herbicide-sensitive control and from each other (p<0.0001), indicating dicamba resistance in M32 is not as high as in 9425. Nevertheless, the ED50 estimate for both resistant populations were greater than the recommended field rate of dicamba (560g ha^-1^), indicating neither population would be controlled by dicamba in a field setting. Similarly, in the 2,4-D experiment, the ED50 estimate for M32 was 5.0 (p<0.0001) times greater than that of the herbicide-sensitive control and greater than the recommended field rate for 2,4-D (approximately 560 g ha^-1^). It is worth noting that the ED50 estimate for the herbicide-sensitive population (543 g ha^-1^) is nearly equal to the recommended field rate for 2,4-D application. Thus, 2,4-D is not likely to provide good control of kochia in general.

**Figure 1.**
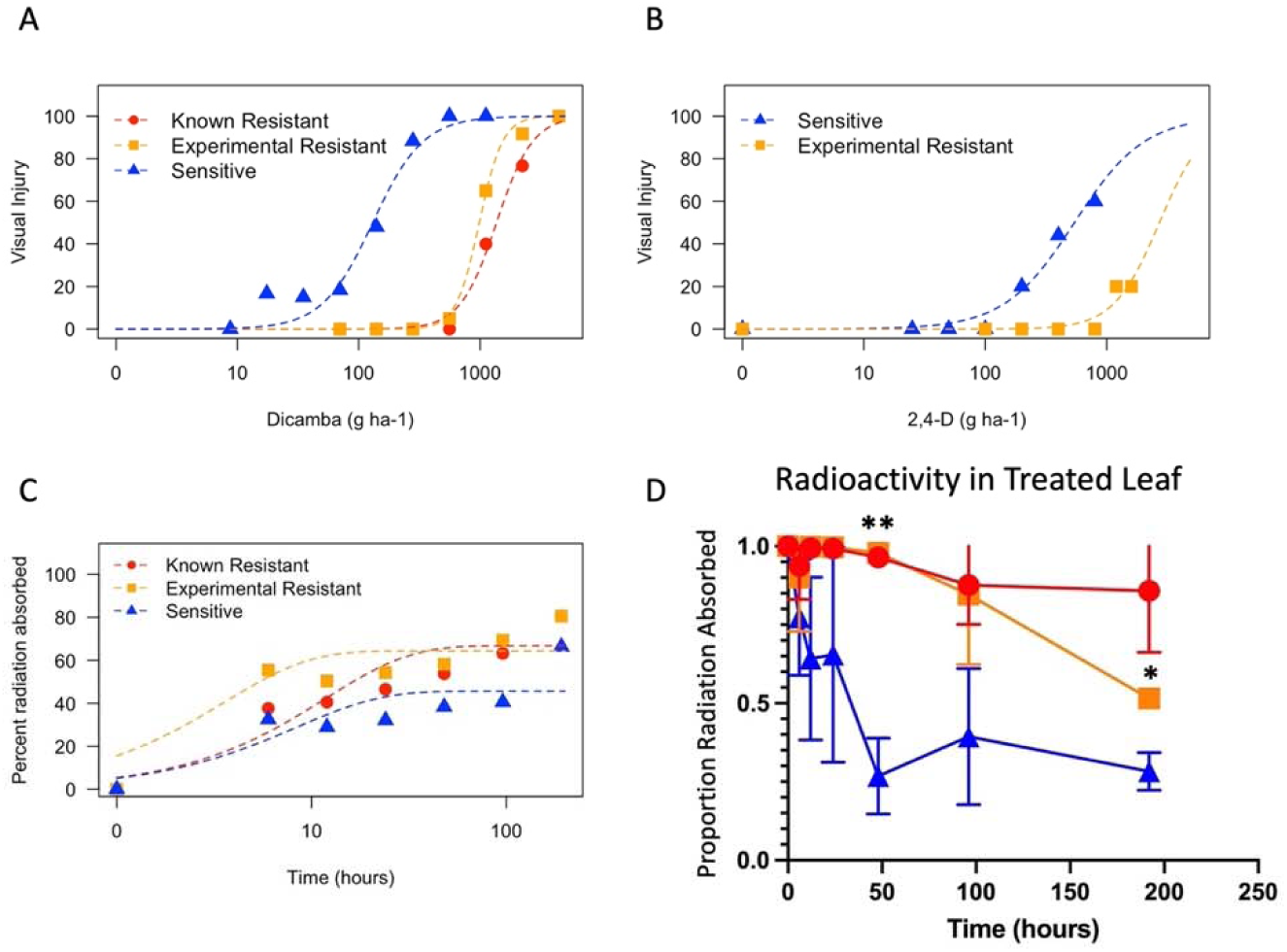
Dose response curves showing magnitude of resistance towards dicamba (A) and 2,4-D (B) for the M32 population of *Bassia scoparia* (Experimental Resistant) compared to a known dicamba-sensitive population and known dicamba-resistant population (LeClere et al., 2018; Pettinga et al., 2018). C) Time course of ^14^C dicamba absorption showing lack of herbicide absorption is not associated with dicamba resistance in dicamba-resistant populations of *Bassia scoparia*. D) Proportion of absorbed ^14^C dicamba retained in the treated leaf showing less dicamba exits the treated leaf in dicamba-resistant populations compared to a dicamba-sensitive population. Error bars represent standard error. *=p<0.05, **=p<0.01. Populations are represented by the same colors and symbols as in A-C.

While herbicide-sensitive plants show epinasty starting approximately 5 days after treatment with dicamba or 2,4-D, resistant plants in our study do not show any symptoms when treated with a field rate of either herbicide. This lack of initial injury suggests that resistant plants either rapidly metabolize/sequester the herbicides before they cause injury (Behrens et al., 2007; Ge et al., 2010) or possess mutant target site(s) that reduce their interaction with their molecular target(s) (Walsh et al., 2006; LeClere et al., 2018; de Figueiredo et al., 2022b). Highly efficient resistance has been engineered into some crops using genes coding for metabolic enzymes from bacteria (Behrens et al., 2007; Wright et al., 2010). In these cases, no injury symptoms are observed at field rates. However, in documented cases of metabolic auxin herbicide resistance, resistant plants often do show symptoms of herbicide damage but eventually recover (de Figueiredo et al., 2022a; Todd et al., 2024).

To understand if physiological mechanisms prevent dicamba from being absorbed or translocated within resistant plants, radiolabeled dicamba was used to track its uptake and movement in dicamba-resistant and -sensitive plants. Significantly less dicamba was absorbed by herbicide-sensitive plants over 192 h compared to the M32 and 9425 populations (p=0.036 and p=0.019 respectively), but the rate of absorption (time required for 90% absorption) was not significantly different between populations (Figure 1C, Table S3). Therefore, reduced dicamba absorption is not responsible for resistance. Time series data for the translocation of radiolabeled dicamba out of the treated leaf did not fit well to any tested models, so t-tests were used to determine if means differed between resistant and sensitive populations at each time point (Figure 1D). Dicamba-resistant plants translocated less dicamba out of the treated leaf compared to the sensitive plants. These results suggest that the dicamba resistance mechanism in the experimental population is caused by a mutation in *BsAUX/IAA16*, which is similar to what has been described by Pettinga et al. (2018) and LeClere et al. (2018) and that is caused by a mutation in *BsAUX/IAA16*.

### A single locus on chromosome 4 is associated with dicamba resistance

A quantitative trait loci (QTL) mapping approach was taken to identify regions of the genome associated with dicamba resistance. Segregating F_3_ populations were developed from multiple independent biparental crosses using a dicamba-resistant plant as the male parent and a dicamba-sensitive plant as the female parent. We expected to find a single dominant locus controlling dicamba-resistance because the ratio of alive:dead F_3_ plants following dicamba treatment at a rate of 560 g ha^-1^ (200:85) did not significantly differ from 3:1 (p = 0.06). We hypothesize that the over-abundance of dead (herbicide-sensitive) plants may be caused by a fitness penalty associated with the dicamba resistance mechanism that reduces germination frequency (see section on fitness cost below). A novel double-digest restriction site associated DNA sequencing (ddRADseq) protocol was used to determine genotypes of 103 segregating F_3_ plants for 1,592 variant loci across the genome. A genome scan associating this genotype information to visual injury rating identified one significant QTL that peaks at 88 Mbp on chromosome 4 (Figure 2A). This interval contains *BsAUX/IAA16* at 87.4 Mbp, a gene that is important for auxin perception and has been implicated in auxin herbicide resistance in kochia and a closely related species (LeClere et al., 2018; Ghanizadeh et al., 2024). In plants from an independent segregating F_3_ line, the genotype of *BsAUX/IAA16* explained 43% of variation in visual injury following dicamba treatment at 560 g ha^-1^ (Figure 2B). While survival at this rate of dicamba may be dominant, visual injury is incompletely dominant (Figure 2B) with a calculated degree of dominance of −0.58 (Falconer, 1964). This dominance has important implications for the spread of herbicide resistance alleles. For instance, a dominant trait is likely to spread more quickly throughout a population, but is less likely to be driven to fixation under selection (Holmes et al., 2022).

**Figure 2.**
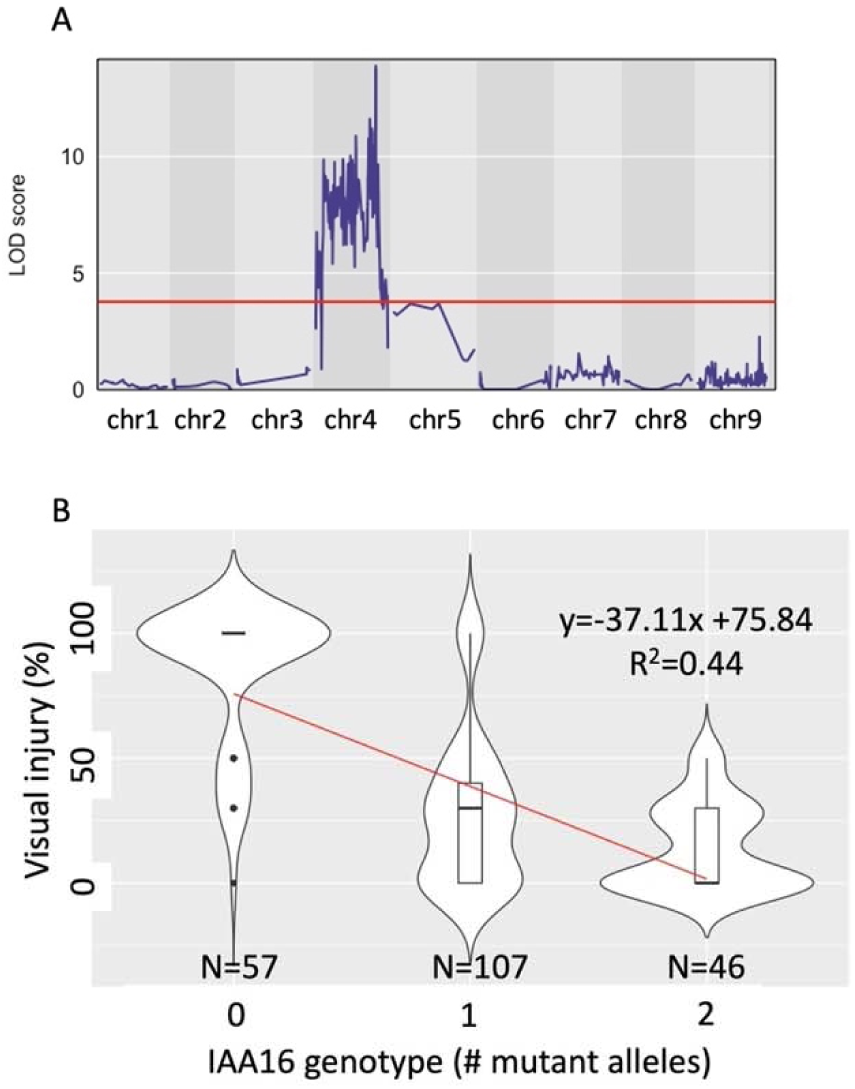
A) Quantitative trait loci (QTL) genome scan from a segregating F_3_ population derived from a biparental cross of herbicide-resistant and -sensitive *Bassia scoparia* plants. Red line represents a significance threshold determined by Bonferroni-adjustment of alpha=0.05 followed by conversion to LOD as described by Nyholt (2000). B) Violin plot displaying the effect of *BsAUX/IAA16* genotype on visual injury 21 days after treatment with dicamba in an independent segregating F_3_ population of *Bassia scoparia*. Red line represents a linear model fitted to all points with equation listed in black. Number of observations for each genotype listed at the base of the graph.

### A transposable element insertion in BsIAA16 causes a change in exon splicing

Sequencing transcripts of *BsAUX/IAA16* from plants homozygous for either parental allele of the QTL discovered on chromosome 4 revealed near identical translated protein sequences except for the region around the degron domain (Figure 3B; Table S4). The gene model for *BsAUX/IAA16* predicts 5 exons in wild type plants with the degron of *BsAUX/IAA16* located near the splice junction of exons one and two (Figure 3A). To understand the sequence of genomic DNA in and around *BsAUX/IAA16*, a plant homozygous for the resistant parent allele of *BsAUX/IAA16* was used for PacBio HiFi sequencing. After *de novo* assembly, we discovered a 3,466 bp insertion at the beginning of exon two of *BsAUX/IAA16* (Figure 3A). The insertion disrupts splicing of *BsAUX/IAA16*, replacing 16 base-pairs at the beginning of exon two in the wildtype with the final 19 bases of the insertion in the mutant (Figure 3A). The mutant allele will herein be referred to as *BsAUX/IAA16_Mut_*, while the wildtype allele will be referred to as *BsAUX/IAA16_WT_*. This insertion results in the addition of one amino acid and the replacement of five more in the final protein product (Figure 3B; Table S4). The new splice sites contain canonical GT and AG motifs. This change includes a substitution of a glycine to threonine at the N-terminal end of degron domain (Figure 3B; Table S4). LeClere et al. (2018) showed that a substitution of this glycine to asparagine in BsAUX/IAA16 is sufficient for dicamba resistance in kochia. Similarly, Ghanizadeh et al. (2024) associate a substitution of this glycine to aspartic acid with dicamba resistance in *Chenopodium album*, a closely related species. Finally, NMR experiments by Ramans-Harborough et al. (2023) show that this glycine is involved in the interaction between AUX/IAA and TIR1. Given the demonstrated importance of this specific glycine residue, we hypothesize that the glycine to threonine substitution is the major driver of dicamba resistance. Protein modeling shows that the larger side group of threonine contacts the surface of the TIR1 binding pocket and causes steric hindrance of the IAA/TIR1 interaction (Figure 3C). The rest of the mutant residues do not interact with the binding pocket of TIR1 and are within a disordered domain of AUX/IAA; thus, they are unlikely to affect IAA/TIR1 interactions (Tan et al., 2007; Ramans-Harborough et al., 2023).

**Figure 3.**
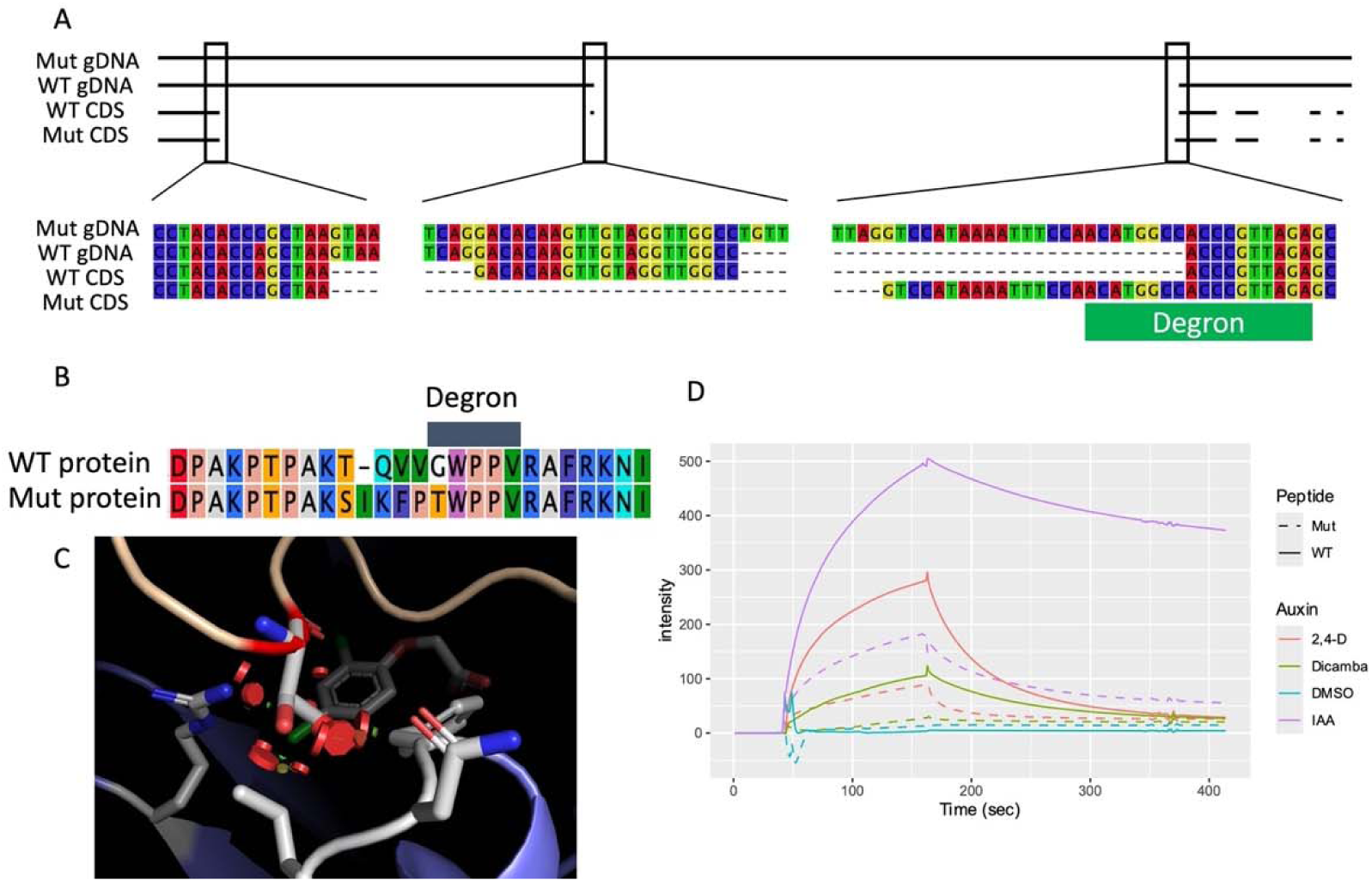
Characterization of a transposable element insertion in *BsAUX/IAA16* and its effect on protein structure and function. A) Genomic and coding sequence for *BsAUX/IAA16* alleles detected in a mapping population segregating for dicamba resistance. B) Amino acid sequence of mutant (Mut) and wildtype (WT) isoforms of BsAUX/IAA16 around the degron domain. C) Crystal structure of IAA7 (gold) bound to TIR1 (blue) showing the glycine 127 residue (red) superimposed over the predicted structure if this residue is substituted for threonine, colored by element (carbon=grey, oxygen=red, nitrogen=blue, chlorine=green). Residues of TIR1 that are within 4Å of the threonine residue are colored by element, and 2,4-D is in the background, colored by element. Steric clashes are shown as red discs. D) Binding/dissociation of BsIAA16 peptides with AtTIR1 in the presence of various auxins as determined by surface plasmon resonance experiments.

The retrotransposon insertion in *BsAUX/IAA16_Mut_* has 5 bp target site duplications flanking 429 bp long terminal repeats (LTRs), identifying it as a Class I LTR element. No protein coding sequences were found within this insertion, suggesting this element is non-autonomous and relies on another transposable element for transposition. In order to search for the corresponding autonomous element, we aligned the insertion to the *de novo* assembly built from a dicamba-resistant plant and the reference genome built from a herbicide-sensitive plant. This identified a near perfect match (99.9% identity) 1 Mbp away on chromosome 4 and another match (96.7% identity) on chromosome 2 in both assemblies (Figure 4B, Table S5). Because the site on chromosome 4 is the most similar across the genome and retrotransposons use a “copy and paste” mechanism of transposition, we hypothesize this to be the donor location for the retrotransposon insertion into in *BsAUX/IAA16_Mut_*. A maximum likelihood phylogenetic tree supports this idea, as the sequences from both assemblies form a distinct clade from those on chromosome 2 (Figure S2). When comparing the insertion in *BsAUX/IAA16_Mut_* with its donor site on chromosome 4, there is one single nucleotide polymorphism (SNP) and two single base pair insertions (Table S5). These insertions are found in homopolymer runs of at least 9 bp and are likely artifacts of the sequencing technology (Espinosa et al., 2024). This similarity suggests very recent transposition. In fact, the 429 bp LTR sequences of the insertion in *AUX/IAA16* are identical. Given the approximated rate of mutation in *Arabidopsis thaliana* of 7×10^−9^ base substitutions per site per generation (Ossowski et al., 2010), we estimate a new single base substitution to be generated in either one of these 429 bp regions approximately every 166,500 generations. This estimation is very crude, but it suggests that the insertion is extremely young in the context of evolutionary time. Future research using collections of kochia in its native range could test whether this adaptation is the consequence of selection on standing genetic variation or de novo genetic diversity that was generated in the time of modern agriculture (Kreiner et al., 2022).

**Figure 4.**
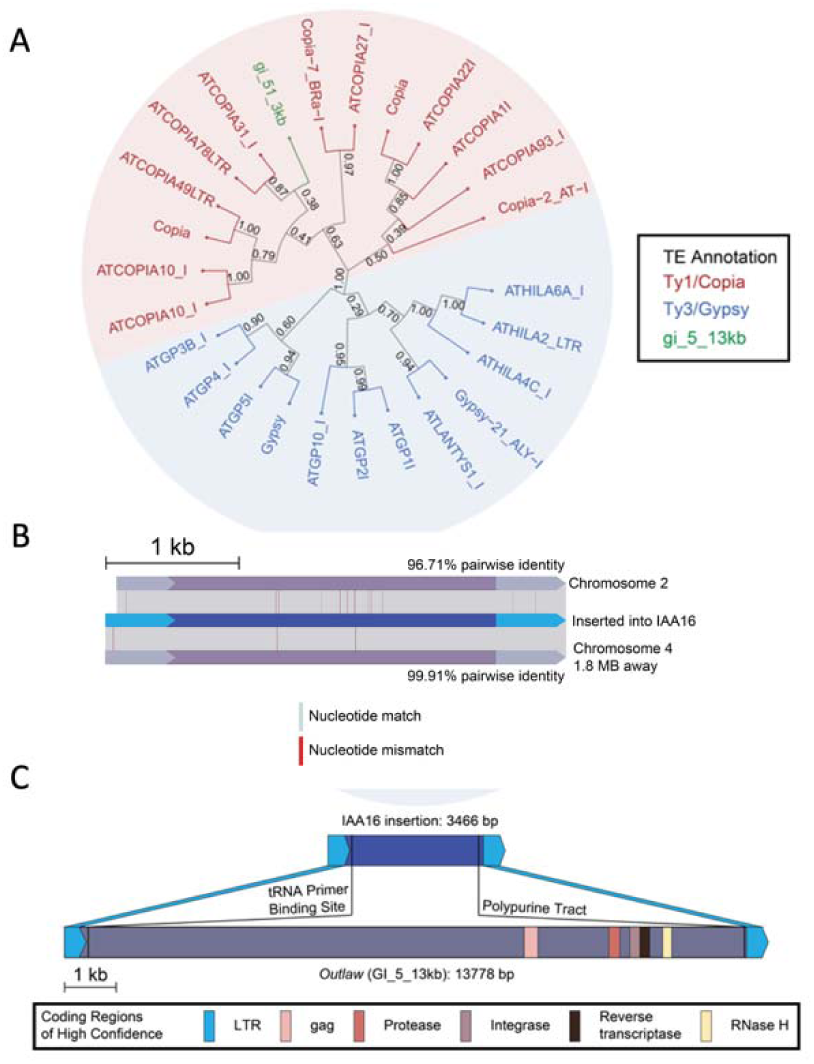
Characterization of autonomous element driving transposition of the retrotransposon that was inserted in *BsAUX/IAA16*. A) Maximum likelihood phylogenetic tree showing evolutionary relationships between autonomous LTR elements from Arabidopsis and the autonomous *Outlaw* element discovered in *Bassia scoparia*. B) Alignment of *Outlaw* elements similar to the one inserted in *BsAUX/IAA16*. C) Comparison of nonautonomous (top) and autonomous (bottom) versions of *Outlaw*. Colored blocks represent regions necessary for retrotransposition.

Since we had identified a likely donor retrotransposon and had not detected any protein coding sequences within this element, there must have been an autonomous element that retrotransposed the donor from chromosome 4 into *BsAUX/IAA16*. To identify the autonomous version(s) of the retrotransposon within *BsAUX/IAA16*, BLAST was first used to find similar LTRs in the previously assembled reference genome (Hall et al., 2023). Putative LTRs were first filtered by sequence similarity and distance between other putative LTRs. Matches were only considered if the putative LTRs were in the same orientation. Once matches were found, identification of required retrotransposon domains was performed for each match. This process led to the identification of a single autonomous element that contained all domains required for transposition (Table S6). This 13 kb putative autonomous retrotransposon, named gi_5_13kb, was found on chromosome 5 (Figure 4C). There are two known superfamilies of retrotransposons, Ty1/Copia and Ty3/Gypsy. To determine which gi_5_13kb belongs to, we first built a curated list of high-confidence LTR transposons in Arabidopsis which contained all required domains. Then, we used the predicted amino acid sequence of these high-confidence LTR transposons to build a phylogenetic tree with the predicted amino-acid sequence of gi_5_13kb. This element clustered within the Ty1/Copia branch of the tree (Figure 4A). We have subsequently named this novel Ty1/Copia element *Outlaw*. This one protein-coding version of the *Outlaw* seems to be driving retrotransposition of the non-autonomous element that generated herbicide resistance in this population of kochia.

### BsIAA16_Mut_ expression is sufficient for dicamba resistance in Arabidopsis thaliana

To demonstrate the role of *BsAUX/IAA16_Mut_* in dicamba resistance, stable transgenic lines of *Arabidopsis thaliana* expressing *BsAUX/IAA16_WT_* or *BsAUX/IAA16_Mut_* were generated through the floral dip method. When seeds were grown on phytoagar plates without herbicide, Arabidopsis expressing *BsAUX/IAA16_Mut_* grew shorter roots compared to those expressing *BsAUX/IAA16_WT_* or with no transgene (Figure 5A). This is evidence of a likely fitness cost associated with disrupting normal auxin signaling in roots due to mutant Aux/IAA over-expression as seen by de Figueiredo et al. (2022b) and LeClere et al. (2018). When grown on media containing 5 μM dicamba, only seedlings expressing *BsAUX/IAA16_Mut_* were able to grow and had significantly longer roots than seedlings expressing *BsAUX/IAA16_WT_*or no transgene when normalized to the length of roots grown on solvent-only media (Figure 5B). A similar result was observed when seedlings were grown on media containing 0.5 μM IAA (Figure 5C), suggesting this variant of AUX/IAA16 also reduces the perception of natural auxin. No seeds grew roots on media containing 5 μM or 0.5 μM 2,4-D.

**Figure 5.**
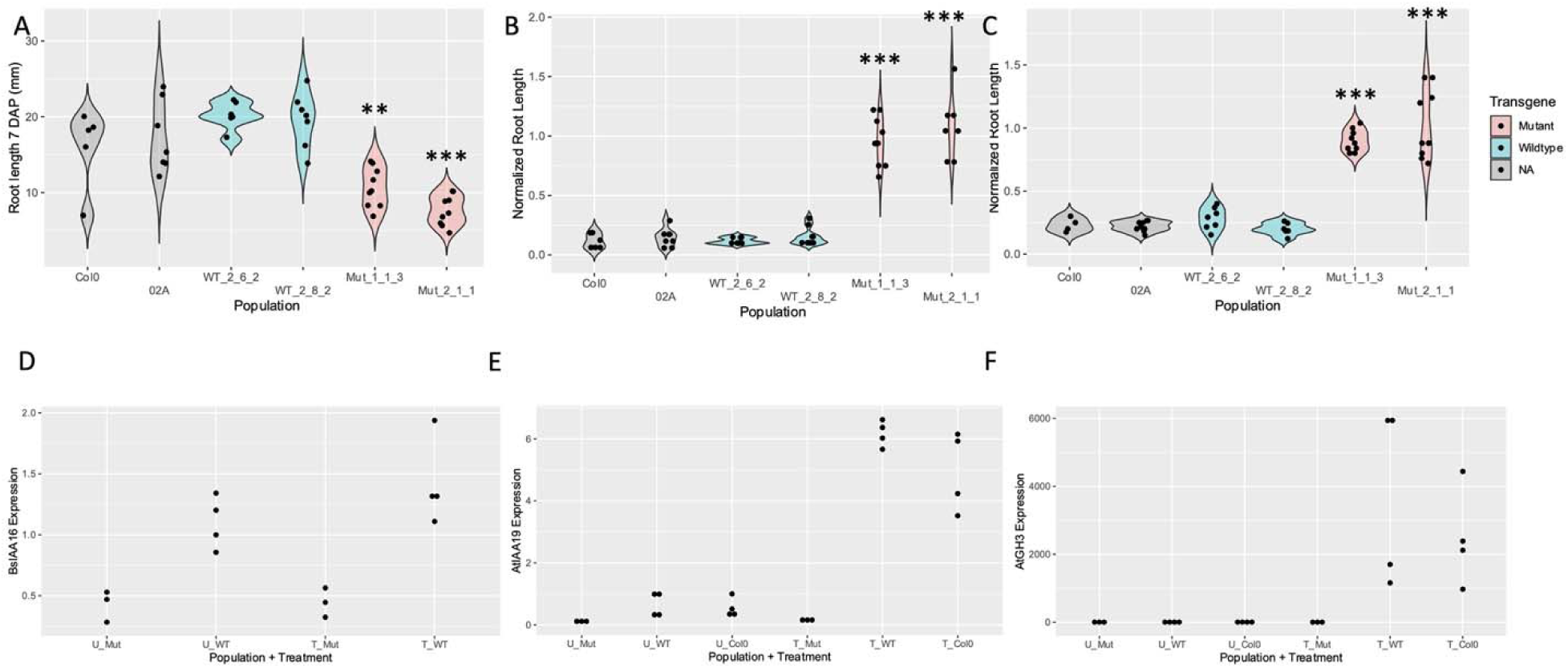
Evaluation of *BsAUX/IAA16* alleles in *Arabidopsis thaliana*. A) Root length of plants expressing either *BsAUX/IAA16_Mut_* (red), *BsIAA16_WT_* (blue), or no transgene (grey) grown on phytoagar media. Proportional root length (compared to ethanol-only control) of the same populations when grown on media containing 5 μM dicamba (B) or 0.5 μM IAA (C). P values in A-C are from t-tests comparing means to the Col0 population. Relative expression of the *BsAUX/IAA16* transgene (D) and the auxin response genes *AtAUX/IAA19* (E) and *AtGH3.3* (F) in plants expressing *BsAUX/IAA16_Mut_* (Mut), *BsAUX/IAA16_WT_*(WT), or no transgene (Col0) that were either treated (T) or untreated (U) with 140 g dicamba ha^-1^ six hours after treatment. Each observation is represented by a black dot. **=p<0.01, ***=p<0.001.

To confirm the results from root growth assays that indicate *BsAUX/IAA16_Mut_* is sufficient for dicamba resistance, plants of select T_3_ lines either expressing *BsAUX/IAA16_WT_*, *BsAUX/IAA16_Mut_*, or no transgene were grown in the greenhouse and treated with 140 g dicamba ha^-1^. Only one of the lines expressing *BsAUX/IAA16_Mut_* showed less visual injury than the other lines tested and appeared stunted and darker green than the other lines even when no dicamba treatment was administered (Figure S3). Quantitative PCR indicated similar expression levels of *BsAUX/IAA16* between lines expressing *BsAUX/IAA16_WT_* or *BsAUX/IAA16_Mut_* (Figure 5D). However, plants expressing *BsAUX/IAA16_Mut_* had no detected induction of auxin response genes *AtIAA19* or *AtGH3.3* six hours after dicamba treatment (Figure 5E and 5F). These genes are known to have increased expression following exposure to natural or synthetic auxins (Tatematsu et al., 2004; Gleason et al., 2011; Pettinga et al., 2018; de Figueiredo et al., 2022b).

### Reduced binding of BsAUX/IAA16_Mut_ to auxin receptor protein TIR1 provides a molecular explanation of resistance

To provide a molecular explanation of the role of *BsAUX/IAA16_Mut_*in dicamba resistance, we measured binding of the Arabidopsis auxin receptor protein AtTIR1 to the WT and mutant BsAUX/IAA16 degron sequences using surface plasmon resonance (SPR; de Figueiredo et al., 2022b; Prusinska et al., 2023). Binding was auxin-dependent. In the presence of indole-3 acetic acid (IAA), binding of the WT peptide was strong and persistent (dissociation of the co-receptor complex was slow; Fig3D). Binding of the WT peptide in the presence of 2,4-D and dicamba was somewhat lower than with IAA as reported previously (Prusinska et al., 2023) with binding amplitudes at the end of the binding phase 56% and 21% of the IAA value, respectively. Binding to the mutant degron peptide was poorer in all respects. The binding amplitudes were lower than for the WT degron (IAA = 37%, 2,4-D 18% and dicamba 5%), and the complexes dissociated more rapidly. In summary, the assembly of the co-receptor complex with the mutant degron was greatly reduced, and when formed, its lifetime was shorter. This would likely lead to far less ubiquitination, higher concentrations of BsAux/IAA16_Mut_ remaining in the cell and reduced auxin signal strength. The greatly reduced signals in the presence of the synthetic auxins 2,4-D and dicamba probably account for resistance as recorded in this line.

### The BsIAA16_Mut_ allele is associated with a reduction in fitness

Because we observe lower binding of *BsAUX/IAA16_Mut_* with TIR1 in the presence of natural auxin, we expect this change to also confer a fitness penalty in the absence of synthetic auxin application (LeClere et al., 2018; Wu et al., 2021; de Figueiredo et al., 2022b). To formally test this, kochia plants were grown from an F_3_ family that is segregating for the *BsAUX/IAA16* locus and fitness traits were measured without the application of auxin. The ratio of genotypes significantly differed from the expected 1:2:1 (p<0.0001), with fewer than expected plants with the *BsAUX/IAA16_Mut_*allele (Figure 6). All seedlings that emerged were used in the experiment, so we hypothesize that *BsAUX/IAA16_Mut_* causes reduced germination. Further, *BsAUX/IAA16_Mut_* was associated with shorter plants, both at transplanting and at maturity (Figures 6A and 6B).

**Figure 6.**
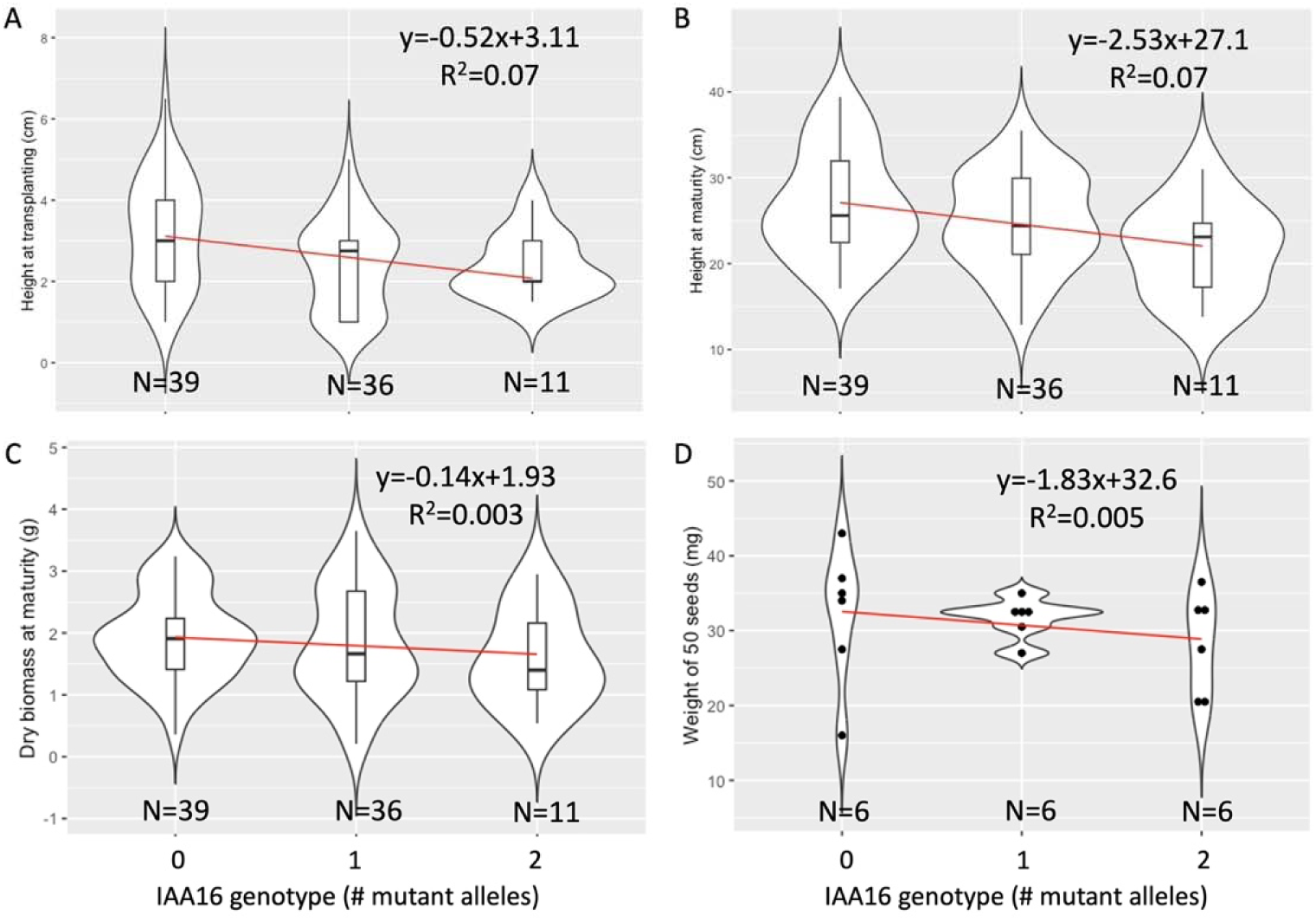
Effect of *BsAUX/IAA16* genotype on plant height at transplanting (A) or maturity (B), dry biomass accumulation at maturity (C), or weight of 50 seeds (D) for kochia plants from an F_3_ population segregating for dicamba resistance. Number of mutant alleles indicate homozygous wildtype (0), heterozygous (1), or homozygous mutant (2). Linear models fitted to the data are plotted in red with equations listed at the top right of each pane. Replicate number is listed at the base of each violin with either boxplots or data plotted as dots within each violin.

However, *BsAUX/IAA16* did not have a significant effect on above ground biomass (Figure 6C) or average seed weight (Figure 6D). Thus, while the *BsAUX/IAA16_Mut_*allele is associated with reduced plant height, plants with this allele accumulate similar biomass and produce similarly sized seeds. Taken together, these results suggest this herbicide resistance mechanism does slightly reduce plant fitness under normal (non-selective) conditions, but not enough to cause it to be completely purged from the population. Nevertheless, evidence of a fitness cost associated with herbicide resistance has implications for kochia management. For instance, if dicamba is not used for several years, the frequency of *BsAUX/IAA16_Mut_* is expected to decrease. Future experiments to investigate allele frequency across generations under competition or abiotic stress will help clarify the real-world ecological consequences of the *BsAUX/IAA16_Mut_* allele (Wu et al., 2018). Hopefully, the negative pleotropic effect of *BsAUX/IAA16_Mut_* may help limit the spread of this resistance allele and allow producers to better manage resistant populations that have already established. This will be especially important as high gene flow in kochia has facilitated rapid spread of other herbicide resistance alleles and mitigated negative effects of genetic bottlenecking following selection for resistant phenotypes (Martin et al., 2020; Ravet et al., 2021).

Taken together, our results further emphasize the importance of the glycine residue within the degron domain of AUX/IAA proteins in auxin signaling. They also reinforce the importance of *BsAUX/IAA16*, specifically, in the action of synthetic auxin herbicides. We describe the first instance of the insertion of a mobile genetic element into the coding sequence of a gene to create an herbicide resistance allele through gain-of-function. The genome of kochia may be especially active for transposable element activity, as the insertion of a mobile genetic element is also associated with gene duplication resulting in glyphosate resistance in this species (Patterson et al., 2019; Ravet et al., 2021; Hall et al., 2023). These results highlight the amazing adaptability of weedy genomes, and the remarkable ways genomes evolve in the face of strong selection pressures (Montgomery et al., 2024).

## Experimental Procedures

### Plant growth and herbicide treatment conditions

Unless otherwise noted, all plants were grown in greenhouses at Colorado State University in 3.8-cm by 3.8-cm by 5.8-cm pots containing fine-grade potting mix (Fafard #2-SV; American Clay Works, Denver, CO). Temperatures were held between 22C and 24C, and supplemental light was provided by liquid halogen grow lights to ensure a photoperiod of 14h/10h. Herbicide applications were made using a moving overhead single-nozzle sprayer (DeVries Manufacturing, Hollandale, MN) calibrated to deliver 187 L ha^−1^. Engenia (BASF,

Research Triangle Park, NC) was used for dicamba applications, and Clean Amine (Loveland Products, Inc., Loveland, CO) was used for 2,4-D applications. For kochia, herbicide applications took place when plants were 10-15 cm tall, and for Arabidopsis, plants were sprayed when they had 10-12 leaves and before bolting.

### Plant material

The M32 population of kochia was collected near Akron, Colorado as part of an herbicide resistance survey conducted between 2012 and 2014 (Westra et al., 2019). Seeds from this field collection were grown, and the resulting plants were treated with 560 g dicamba ha^-1^. Survivors were open-pollinated, and their seed was combined to form a composite dicamba-resistant seed lot. Approximately 50 seeds from this combined lot were grown and treated with 560 dicamba ha^-1^. The individual with the least herbicide injury was used as a male parent in a biparental cross with an individual from the population 7710, an inbred, herbicide-susceptible population (Preston et al., 2009) that was used to generate the reference genome assembly for kochia (Patterson et al., 2019; Hall et al., 2023). To make the cross, immature flowers of the dicamba-sensitive plant were emasculated using forceps under a microscope, and the emasculated flowers were labeled. The two plants were then grown to maturity together in a pollen-exclusion tent.

Seeds from the dicamba-sensitive plant were collected and grown; the resulting plants were treated with 560 g dicamba ha^-1^. A Kompetative Allele Specific PCR (KASP) assay that detects *BsAUX/IAA16_Mut_* was used to confirm survivors as hybrids (primer sequence and thermocycler conditions found in Table S7). Several confirmed hybrid plants were grown in pollen exclusion tents and self-pollinated to create several F_2_ families. Plants from two F_2_ families were grown, but because of low seed production in the F_1_ generation, plants were allowed to self-pollinate with no selection to produce F_3_ families.

### Dose response

Plants were grown from the 7710 (Preston et al., 2009) and 9425 (LeClere et al., 2018) populations as well as from the M32 composite resistant seed lot. Plants were treated with one of several rates of dicamba. These rates included 0, 8.75, 17.5, 35, 70, 140, 280, 560, and 1120 g dicamba ha^-1^ for the herbicide-sensitive population and 0, 70, 140, 280, 560, 1120, 2240, and 4480 g dicamba ha^-1^ for the herbicide-resistant populations. Six uniform plants were used for each population at each herbicide rate before being returned to the greenhouse. Visual injury and survival were rated 21 days after treatment (DAT). The drc package v3.0-1 (Ritz et al., 2015) in R v4.0.2 (R-Core-Team, 2021) was used to fit the injury data to a 2-parameter log-logistic dose response curve. Significant difference is in the ED50 parameter were determined using the compParm function from the drc package (Ritz et al., 2015).

### Herbicide absorption and translocation profiles

Kochia plants from M32, 7710, and 9425 were germinated in a growth chamber (60% relative humidity, 21/18 C, and 16/8 h photoperiod) in potting soil and transplanted to fine sand when they reached ∼3 true leaves. The plants were irrigated with fertilizer until the plants reached 10 cm in height. At this point, aluminum foil was used to cover the second youngest fully expanded leaf while the plants were sprayed with 560 g dicamba ha^-1^. After spraying, the aluminum foil was removed, the covered leaf was then marked and treated with 10 μL of a ^14^C-labeled dicamba solution (total radioactivity of 3.33 KBq or 200,000 dpm per plant). Plants were returned to the growth chamber until sampling at 3, 6, 12, 24, 48, 96, and 192 h after treatment. The treated leaf, remaining above ground tissue, and below ground tissue of three biological replicates were separated for each population at each time point. The treated leaf was washed in 10 mL 10% methanol + 1% NIS, and radioactivity in this wash solution was quantified in 10 mL of scintillation mixture (Ecoscint XR, National Diagnostics, Atlanta, GA) using liquid scintillation spectrometry (Packard Tricarb 2300TR, Packard Instrument Co., Meriden, CT).

Plant tissue was dried in an oven at 60 C for at least 14 d before oxidation in a biological oxidizer (OX500; RJ Harvey Instrument Co., Tappan, NY) followed by radioactivity measurement by liquid scintillation spectrometry. Absorption time series data was fitted to a rectangular hyperbolic model, and the parameters “absorption max” (*Amax*) and “time to 90 percent absorption” (*t90*) were estimated and compared in R (Kniss et al., 2011; Ritz et al., 2015; R-Core-Team, 2021). Because the time series data for translocation of dicamba outside of the treated leaf did not fit any of the models proposed by Kniss et al. (2011), t-tests were used to determine if the amount of dicamba retained in the treated leaf was different between the M32 and 7710 populations at each time point.

### QTL mapping

Plants from several F_3_ families derived from a biparental cross described above were grown and treated with 560 g dicamba ha^-1^. Two families that segregated for survival following dicamba treatment (4-1-1 and 4-5-10, each from a different F_1_ hybridization event) were used for QTL mapping. Approximately 150 plants from the F_3_ family 4-5-10 were grown, and a single young leaf was sampled and frozen in liquid nitrogen before treatment with 560 g dicamba ha^-1^. Visual injury on a scale from 0-100 was rated for each plant 21 DAT, and DNA was extracted from each plant (including the original parents of the cross) using the CTAB method (Doyle & Doyle, 1990). The concentration of each DNA sample was quantified using the Qubit™ dsDNA BR kit (Thermo Scientific, Waltham, MA) and diluted to 20 ng/μl. A double-digest restriction-site associated DNA sequencing (ddRADseq) protocol was developed in conjunction with the University of Minnesota Genomics Center (Minneapolis, MN) by estimating the number of RAD sites across the reference genome (Patterson et al., 2019) for several enzyme combinations and then empirically testing the BtgI-TaqI combination on several samples at multiple sequencing depths. DNA from the two original parents and 103 plants from the 4-5-10 F_3_ family were used for ddRADseq with BtgI and TaqI as the restriction enzymes with a target of 4 million 150 bp single end reads per sample. Library prep and sequencing was completed by the University of Minnesota Genomics Center.

Reads from each sample were passed through a variant calling pipeline that can be found at github.com/JMont12/dicamba_kochia. Briefly, raw reads were trimmed with Trimmomatic v0.36 (Bolger et al., 2014) and aligned to the reference genome of kochia (Hall et al., 2023) using the burrows-wheeler aligner v0.7.17 (Li & Durbin, 2009). The alignments were processed with samtools v1.15 (Li et al., 2009) and passed to GATK v4.2.0 (Poplin et al., 2018; Van der Auwera & O’Connor, 2020) to call variants and filter them based on depth, quality, and strand bias. Variants were further filtered using custom scripts to only include biallelic single nucleotide polymorphisms that were homozygous and different between the two original parents. Filtered variants and phenotype data were used to conduct a genome scan within the qtl2 package in R v0.32 (Broman et al., 2019). Because a permutation test for significance was not possible, a significance threshold was established through a Bonferroni correction of alpha=0.05 considering the number of tests to equal the number of filtered markers used in the scan then converting the adjusted p-value to LOD as described by Nyholt (2000). The genotype around the QTL discovered on chr4 was determined for 210 plants from another F_3_ family (4-1-1) using the KASP assay developed for the *BsAUX/IAA16* gene (primer sequence and thermocycler conditions in Table S6). A simple linear model was fitted using visual injury 21 DAT as the dependent variable and *BsAUX/IAA16* genotype as the independent variable and plotted using ggplot2 v3.4.4 (Wickham, 2016) in R (R-Core-Team, 2021).

Similarly, to quantify the effect of *BsAUX/IAA16* genotype on fitness traits, ∼300 seeds of an F3 family segregating for the *BsAUX/IAA16* locus were sown into damp soil. Three weeks after planting, all emerged seedlings were transplanted into individual pots (10.5 x 10.5 x 12 cm) filled with potting soil and irrigated evenly throughout their lifecycle. Plants were spaced evenly, watered sufficiently, and not fertilized to reduce competition and environmental variation. Plant height was measured at transplanting and maturity. After allowing the plants to senesce naturally, above ground biomass was collected and weighed. For a subset of plants, seeds were harvested and a random sample of 50 seeds was used to determine average seed weight for each sample. During development, a small amount of leaf tissue was collected from each plant and used to determine the genotype of the *BsAUX/IAA16* locus using the KASP assay described above. Genotype and phenotype information was used to fit linear models and plotted using ggplot in R.

### BsIAA16_M32_ sequencing

Total RNA was extracted using the Direct-zol RNA Microprep kit (Zymo Research, Irvine, CA) from individuals of the 4-5-10 selected line that were homozygous for either parental allele of *BsAUX/IAA16* based on the results of KASP testing. This RNA was used to generate cDNA libraries for each sample using the ProtoScript® II First Strand cDNA Synthesis Kit (New England Biolabs, Ipswich, MA). Primers that bind to the beginning and end of the *BsAUX/IAA16* gene (Table S7) were used with the EconoTaq PLUS Green kit (LGC, Teddington, Middlesex, UK) to amplify the full-length coding sequence. The product of each PCR reaction was run on a 1% agarose gel to ensure a single product, then sent to Azenta (Burlington, MA) for Sanger sequencing using the forward and reverse primers.

To determine the genomic sequence around the *BsAUX/IAA16_Mut_*, 2 g of dark-treated leaf tissue from a plant that was homozygous for *BsAUX/IAA16_Mut_* was sampled and flash frozen in liquid nitrogen. This tissue was sent to Corteva Agriscience Center for Genome Excellence (Johnston, IA) for whole genome PacBio HiFi sequencing to a depth of ∼30X. The resulting reads were assembled with HiFiasm v0.19.5 (Cheng et al., 2021), and *BsAUX/IAA16* was located by BLAST v2.6.0 (Camacho et al., 2009).

### Characterization of transposable element insertion

To find similar elements to the one inserted in *BsAUX/IAA16_Mut_*, the sequence of the 429 bp LTRs of the insertion were aligned to the kochia reference genome assembly (Hall et al., 2023) and to the newly created assembly described above using BLAST (Camacho et al., 2009). Sequential filtering was used to identify the autonomous version of the element. First, alignments of >85% identity and >100bp alignment length were identified as putative LTRs. Next, only putative LTRs that were between 25 kb and 4 kb apart and in the direct orientation were considered potential *Outlaw* transposon family members. Finally, of the elements of the proper length and LTR orientation, we then only considered putative elements that contained all required retrotransposon domains as potential autonomous elements. Necessary retrotransposon domains were identified manually and with TESorter (Zhang et al., 2022). Specifically, the required domains that we used were as follows: gag, pol (RNAseH, Integrase, Protease, Reverse Transcriptase), tRNA Primer Binding Site, and Polypurine Tract.

### Autonomous transposable element characterization

To classify the retro element into one of two possible superfamilies (Ty1/Copia or Ty3/Gypsy), we used the *Arabidopsis thaliana* transposable element annotation dataset from RepetDBv2 (Amselem et al., 2019). We procured all Ty1/Copia and Ty3/Gypsy nucleotide elements and predicted the protein domain sequences from this dataset using TESorter (Zhang et al., 2022). We only used TEs that had all required retrotransposon domains (both pol and gag).

We then manually concatenated all pol domain proteins (RT, RNaseH, Protease, Integrase) as a single polypeptide for each transposable element and then used *Outlaw* predicted pol polypeptide sequence to create a phylogenetic tree. MegaX was used to align all of the proteins using MUSCLE to create a maximum likelihood tree with 500 bootstraps (Kumar et al., 2018).

### Development of transgenic Arabidopsis thaliana lines

The coding sequence of *BsAUX/IAA16_Mut_* and *BsAUX/IAA16_WT_*were amplified from cDNA libraries discussed above using the PrimeSTAR MAX DNA Polymerase kit (Takara Bio USA, Inc., San Jose, CA) and primers with tails that complement restriction sites in the expression vector *pFGC5941* (primer sequences and thermocycler conditions in Table S8).

Empty *pFGC5941* was digested with AscI and BamHI (New England Biolabs, Ipswich, MA), and the *BsAUX/IAA16* amplicons were ligated into the digested vector using the In-Fusion Cloning kit (Takara Bio USA, Inc., San Jose, CA). Plasmid sequencing ensured correct assembly of expression vectors (Plasmidsaurus, Eugene, OR). Vectors containing *BsAUX/IAA16_Mut_* or *BsAUX/IAA16_WT_* were transformed into the GV3101 strain of *Agrobacterium tumefaciens*, and subsequently transfected into the Col-0 genotype of *Arabidopsis thaliana* using the floral dip method (Clough & Bent, 1998). Transformants were selected by spraying T_1_ seedlings with BASTA herbicide (BASF, Research Triangle Park, NC) and confirmed with PCR for the transgene (primers and thermocycler conditions found in Table S9). Confirmed transformants were inbred for two generations, and T_3_ populations fixed for the transgene were identified by spraying seedlings with BASTA herbicide and PCR for the transgene.

### Effect of dicamba on transgenic Arabidopsis lines

Seeds of Col-0 and of fixed T_3_ lines of Arabidopsis, each from a different transformation event, were gas sterilized and plated on media plates containing synthetic or natural auxin as described by de Figueiredo et al. (2022b). Plates were incubated in a growth chamber with a yellow light filter to prevent indole-3 acetic acid degradation. Ten seeds were used per population for each treatment. Root growth was measured 7 d after moving plates to growing conditions. T-tests were used to detect significant differences in root length between lines expressing *BsAUX/IAA16_Mut_* compared to the Col-0 control. Results were plotted with ggplot2 in R.

Plants from select T_3_ lines were grown to the 10-leaf stage and either untreated or treated with 140 g dicamba ha^-1^. Four plants of each population were used for each treatment. Photos were taken 21 DAT to illustrate visual injury. Young leaf tissue was collected from treated and untreated plants 6 h after treatment. Total RNA was extracted from these leaf samples using the Direct-zol RNA Microprep kit (Zymo Research, Irvine, CA), and cDNA libraries were generated using the ProtoScript® II First Strand cDNA Synthesis Kit (New England Biolabs, Ipswich, MA). Following the methods of de Figueiredo et al. (2022b), relative expression was quantified for the *BsAUX/IAA16* transgene (primers and thermocycler conditions available in Table S10) and the auxin response genes *AtIAA19* and *AtGH3.3* using *AtCyclophilin* as a reference gene and the 2^−ΔΔCt^ method.

### In silico mutagenesis of IAA variants

The crystal structure of the degron of IAA7 bound to TIR1 in the presence of 2,4-D (2p1n) was downloaded from the RCSB Protein Data Bank (Tan et al., 2007). Mutagenesis of the G127 residue was conducted and visualized in PyMOL.

### Reeceptor protein preparation and Surface Plasmon Resonance measurement of auxin binding

Receptor protein (AtTIR1) was purified and the SPR assays performed following the methods described in Prusinska et al., 2023. The SPR chip (Series S SA) was loaded with our reference negative control degron peptide mIAA7 (mutated Arabidopsis Aux/IAA7); channel 2 with positive control degron peptide IAA7; channel 3 with BsAUX/IAA16_WT_ degron peptide (biotin-AKTQVVGWPPVRAFRKN-amide) and channel 4 with BsAUX/IAA16_Mut_ degron peptide (biotin-AKSIKFPTWPPVRAFRKN-amide). Similar peptide loads were recorded for all four channels. Purified receptor was passed over all four channels simultaneously in parallel in PBS buffer + 0.01% Tween20, with or without test auxin (IAA; 2,4-D; or dicamba) at a concentration of 50 μM. Binding was recorded for 120 s, and dissociation for 180 s. Data were visualized in BiaEvaluation software.

## Data Availability

The RAD sequencing data from the segregating F_3_ line used in the mapping project is available through NCBI, BioSample IDs SAMN42890277-SAMN42890594 and SRA accession numbers SRR30019316-SRR30019633. Phenotypic data can be found in Supplemental Table S11. PacBio HiFi sequencing used to generate the de novo genome assembly is available through NCBI under BioSample ID SAMN43525227 and SRA accession number SRR30575711. All sequence data is associated with NCBI BioProject PRJNA1141446.

## Acknowledgments

This research was supported in part by BASF SE, by the Colorado Wheat Administrative Committee, and by the USDA National Institute of Food and Agriculture, Hatch project COL00783 to the Colorado State University Agricultural Experiment Station. The authors have no conflict of interest to declare.

