## Supplementary material for "A transposable element insertion in *IAA16* disrupts splicing and causes dicamba resistance in *Bassia scoparia*": Figure S2

Supplemental Information

Table S1. Parameter estimates for 2-parameter log-logistic dose response curves for M32, 9425, and 7710 kochia populations in response to dicamba. The equation fitted for each population was $y = \frac{100}{1+b(\log(x)-\log(ED50))}$ where *y* is predicted visual injury (0-100) 21 days after dicamba treatment, *x* is the rate of dicamba in g $ha^{-1}$, *ED50* is the rate of dicamba required to cause 50% visual injury, and *b* is the slope of the curve at x=ED50.

| Parameter | Estimate | Std. Error |
|---|---|---|
| Slope:7710 | -2.9 | 0.3 |
| Slope:M32 | -4.5 | 0.9 |
| Slope:9425 | -2.1 | 0.3 |
| ED50:7710 | 130.6 | 16.9 |
| ED50:M32 | 988.3 | 27.2 |
| ED50:9425 | 1370.4 | 20.9 |

Table S2. Parameter estimates for 2-parameter log-logistic dose response curves for M32 and 7710 kochia populations in response to 2,4-D. The equation fitted for each population was $y = \frac{100}{1+b(\log(x)-\log(ED50))}$ where *y* is predicted visual injury (0-100) 21 days after 2,4-D treatment, *x* is the rate of 2,4-D in g $ha^{-1}$, *ED50* is the rate of 2,4-D required to cause 50% visual injury, and *b* is the slope of the curve at x=ED50.

| Parameter | Estimate | Std. Error |
|---|---|---|
| Slope:7710 | -1.5 | 0.1 |
| Slope:M32 | -2.3 | 0.4 |
| ED50:7710 | 543.1 | 21.2 |
| ED50:M32 | 2720.7 | 307.8 |

Table S3. Parameter estimates for rectangular hyperbolic models fitted to herbicide absorption data for M32, 9425, and 7710 kochia populations. The equation fitted for each population was $y = (Amax \times t)/(0.11 \times t90 + t)$ where $y$ is percent of applied herbicide that was absorbed, $t$ is time after herbicide application (in hours), $t90$ is the time required for 90% herbicide absorption, and $Amax$ is the maximum amount of herbicide absorbed.

| Parameter | Estimate | Std. Error |
|---|---|---|
| Amax:7710 | 66.8 | 5.6 |
| Amax:M32 | 64.4 | 3.9 |
| Amax:9425 | 45.7 | 5.3 |
| t90:7710 | 27.2 | 9.5 |
| t90:M32 | 8.3 | 4.1 |
| t90:9425 | 19.3 | 14.6 |

Table S4.
CLUSTAL multiple protein sequence alignment of 7710 and M32 alleles of AUX/IAA16. The degron domain is bolded.

```
7710  MLSNERDKYTIDFEETELRLGLGLGIGLAGAADGDQLAKNNNGKRGFSETEGDSSVDLKL
M32   MLSNERDKYTIDFEETELRLGLGLGIGLAGAADGDQLAKNNNGKRGFSETEGDSSVDLKL
      ************************************************************
7710  NLSSSTTTTASTTTTNTTATKTTAENVKESKLDKSVNSGVDQKLKEKVASTTADPAKPTP
M32   NLSSSTTTTASTTTTNTTATKTTAENVKESKLDKSVNSGVDQKLKEKVASTTADPAKPTP
      ************************************************************
7710  AKT-QVVGWPPVRAFRKNIVAAHKKTSDDQTDQKASSNAITSAAFVKVSMDGAPYLRKVD
M32   AKSIKFPTWPPVRAFRKNIVAAQKKTSDDQTDQKASSNAITSAAFVKVSMDGAPYLRKVD
      **: :.  **************:*************************************
7710  LKLYKSYQDLSDALGKMFSSFTIGNCGSQGMKDFMNESKLIDLLNGSEYVPTYEDKDGDW
M32   LKLYKSYQDLSDALGKMFSSFTIGNCGSQGMKDFMNESKLIDLLNGSEYVPTYEDKDGDW
      ************************************************************
7710  MLVGDVPWEMFVGSCKRLRIMKGSEAIGLAPRAVEKCKNRS*
M32   MLVGDVPWEMFVGSCKRLRIMKGSEAIGLAPRAVEKCKNRS*
      *****************************************
```

Table S5. CLUSTAL alignment of transposable elements similar to the one in *AUX/IAA16* from the M32 population. Outgroup represents the version on chromosome 2. Donor represents the conserved version found on chromosome 4. IAA_insertion is the M32-specific version inside of *AUX/IAA16*.

```
IAA_insertion      TGTTGGGATTAAACCCTAATTCATATCAAATTAGTCGGTAATAATAGATCTGCAGAAGCA
M32_donor          TGTTGGGATTAAACCCTAATTCATATCAAATTAGTCGGTAATAATAGATCTGCAGAAGCG
7710_donor         TGTTGGGATTAAACCCTAATTCATATCAAATTAGTCGGTAATAATAGATCTGCAGAAGCG
M32_outgroup       T-----------------------------------------------------------
7710_outgroup      T-----------------------------------------------------------
                   *

IAA_insertion      TACCTGAATCCATGACGAAGATCGGCGGTGGGTTTTGATCTTCCAATTCTTTATGGCTCC
M32_donor          TACCTGAATCCATGACGAAGATCGGCGGTGGGTTTTGATCTTCCAATTCTTTATGGCTCC
7710_donor         TACCTGAATCCATGACGAAGATCGGCGGTGGGTTTTGATCTTCCAATTCTTTATGGCTCC
M32_outgroup       ------------------------------------------CCAATTCTTTATGGCTCC
7710_outgroup      ------------------------------------------CCAATTCTTTATGGCTCC
                                                             ******************

IAA_insertion      TTAGGGTTTCTACTGATGATGGGATGTCAGTGAGAATAGGAAAACCATATGAATCGGGGA
M32_donor          TTAGGGTTTCTACTGATGATGGGATGTCAGTGAGAATAGGAAAACCATATGAATCGGGGA
7710_donor         TTAGGGTTTCTACTGATGATGGGATGTCAGTGAGAATAGGAAAACCATATGAATCGGGGA
M32_outgroup       TTAGGGTTTCTACTGATGATGGGATGTCAGTGAGAATAGGAAAACCATATAAATCGGGGA
7710_outgroup      TTAGGGTTTCTACTGATGATGGGATGTCAGTGAGAATAGGAAAACCATATAAATCGGGGA
                   ************************************************** *********

IAA_insertion      CCATAACCCTTAATGTCTATTTATATACATAGACTCCTTCCTAATCCGCCCATCGTGAAT
M32_donor          CCATAACCCTTAATGTCTATTTATATACATAGACTCCTTCCTAATCCGCCCATCGTGAAT
7710_donor         CCATAACCCTTAATGTCTATTTATATACATAGACTCCTTCCTAATCCGCCCATCGTGAAT
M32_outgroup       CCATAACCCTTAATGTCTATTTATATACATAGACTCCTTCCTAATCCGCCCATCGTGAAT
7710_outgroup      CCATAACCCTTAATGTCTATTTATATACATAGACTCCTTCCTA-TCCGCCCATCGTGAAT
                   ******************************************* ****************

IAA_insertion      AAGGAAAGGCCCATCGGTATCTACACAAATAAAAAACTGATCCCACACTCTATAAAAGAC
M32_donor          AAGGAAAGGCCCATCGGTATCTACACAAATAAAAAACTGATCCCACACTCTATAAAAGAC
7710_donor         AAGGAAAGGCCCATCGGTATCTACACAAATAAAAAACTGATCCCACACTCTATAAAAGAC
M32_outgroup       AAGGAAAGGCCCATCGGTATCTACACAAATAAAAAACTGATCCCACACTCTATAAAAGAC
7710_outgroup      AAGGAAAGGCCCATCG-TATCTAC-CAAATAAAAA-CTGATCCCACACTCTATAAAAGAC
                   **************** ******* ********** ************************

IAA_insertion      GTAAATAGGCCCAATAATAATTACTTAATTGGATCACTTTAGTTTTGGGCCACACCGTAT
M32_donor          GTAAATAGGCCCAATAATAATTACTTAATTGGATCACTTTAGTTTTGGGCCACACCGTAT
7710_donor         GTAAATAGGCCCAATAATAATTACTTAATTGGATCACTTTAGTTTTGGGCCACACCGTAT
M32_outgroup       GTAAATAGGCCCAATAATAATTACTTAATTGGATCACTTTAGTTTTGGGCCACACCGTAT
7710_outgroup      GTAAATAGGCCCAATA-TAATTACTTAATTGGATCACTTTAGTTTTGGGCCACACCGTAT
                   **************** *******************************************

IAA_insertion      GATAGCACATAATACAATTATAACTGAATTGCACACGTATTTGTATTTAGGTCCATAAAA
M32_donor          GATAGCACATAATACAATTATAACTGAATTGCACACGTATTTGTATTTAGGTCCATAAAA
7710_donor         GATAGCACATAATACAATTATAACTGAATTGCACACGTATTTGTATTTAGGTCCATAAAA
M32_outgroup       GATAGCACATAATACAATTATAACTGAATTGCACACGTATTTGTATTTAGGTCCATAAAA
7710_outgroup      GATAGCACA--ATACAATTATAACTGAATTGCACACGA--TTGTATTTAGGTCCATAAAA
                   *********  **************************   ********************

IAA_insertion      TTTCCAACAGTCTCCCACTTGGACCAAATACAATTACAACGTGTGTGATACTTGATAATT
M32_donor          TTTCCAACAGTCTCCCACTTGGACCAAATACAATTACAACGTGTGTGATACTTGATAATT
7710_donor         TTTCCAACAGTCTCCCACTTGGACCAAATACAATTACAACGTGTGTGATACTTGATAATT
M32_outgroup       TTTCCAACAGTCTCCCACTTGGACCAAATACAATTACAACGTGTGTGATACTTGATAATT
7710_outgroup      TTTCCAACAGTCTCCCACTTGGACCAAATACAATTACAACGTGTGTGATACTTGATAATT
```

```
                    ************************************************************

IAA_insertion       GCATTATACTTTATAA-CCTTATGAGCTCAAAATTGCTATCAAAATCTCAACGTCTTTAA
M32_donor           GCATTATACTTTATAA-CCTTATGAGCTCAAAATTGCTATCAAAATCTCAACGTCTTTAA
7710_donor          GCATTATACTTTATAA-CCTTATGAGCTCAAAATTGCTATCAAAATCTCAACGTCTTTAA
M32_outgroup        GCATTATACTTTATAA-CCTTATGAGCTCAAAATTGCTATCAAAATCTCAACGTCTTTAA
7710_outgroup       GCATTATACTTTATAAACCTTATGAGCTCAAA-TTGCTATCAAAATCTCAACGTCTTTAA
                    **************** *************** ***************************

IAA_insertion       ACAATTCAGTCCATTAATTACATCAACACAGGATCAAAGAGATCTATGCTACATTTGCCG
M32_donor           ACAATTCAGTCCATTAATTACATCAACACAGGATCAAAGAGATCTATGCTACATTTGCCG
7710_donor          ACAATTCAGTCCATTAATTACATCAACACAGGATCAAAGAGATCTATGCTACATTTGCCG
M32_outgroup        ACAATTCAGTCCATTAATTACATCAACACAGGATCAAAGAGATCTATGCTACATTTGCCG
7710_outgroup       ACAATTCAGTCCATTAATTACC---ACACAGGATCAAAGAGATCTATGCTACATTTGCCG
                    *********************    ***********************************

IAA_insertion       TAACCAGACCCATCAATGGTCACAATGTCAACATAATTAACGACATGAATCAAGCATGGG
M32_donor           TAACCAGACCCATCAATGGTCACAATGTCAACATAATTAACGACATGAATCAAGCATGGG
7710_donor          TAACCAGACCCATCAATGGTCACAATGTCAACATAATTAACGACATGAATCAAGCATGGG
M32_outgroup        TAACCAGACCCATCAATGGTCACAATGTCAACATAATTAACGACATGAATCAAGCATGGG
7710_outgroup       TAACCAGACCCATCAATGGTCACAATGTCAACATAATTAACGACATGAATCA-GCATGGG
                    **************************************************** *******

IAA_insertion       TGTGTAGCATGGAAATTACATACAAATGTGATCCAAGTATGCCTATTTCCAACTGGTCCA
M32_donor           TGTGTAGCATGGAAATTACATACAAATGTGATCCAAGTATGCCTATTTCCAACTGGTCCA
7710_donor          TGTGTAGCATGGAAATTACATACAAATGTGATCCAAGTATGCCTATTTCCAACTGGTCCA
M32_outgroup        TGTGTAGCATGGAAATTACATACAAATGTGATCCAAGTATGCCTATTTCCAACTGGTCCA
7710_outgroup       TGTGTAGCATGGAAATTACATACAAATGTGATCCAAGTATGCCT-TTTCCAACTGGTCCA
                    ******************************************** ***************

IAA_insertion       CTGTAAACTTTGGTAAGATCAACAATATGATCTAATCGAAGAGTAAACCGAACCGAATAC
M32_donor           CTGTAAACTTTGGTAAGATCAACAATATGATCTAATCGAAGAGTAAACCGAACCGAATAC
7710_donor          CTGTAAACTTTGGTAAGATCAACAATATGATCTAATCGAAGAGTAAACCGAACCGAATAC
M32_outgroup        CTGTAAACTTTGGTAAGATCAACAATATGATCTAATCGAAGAGTAAACCGAACCGAATAC
7710_outgroup       CTGTAAACTTTGGTAAGATCAACAATATGATCTAATCGAAGAGTAA-CCGAACCGAATAC
                    ********************************************** *************

IAA_insertion       CTTATTTCTGCAGAAAATACCTTAAACCTTAATATCCGAATTAGAACTTCATCATACAAG
M32_donor           CTTATTTCTGCAGAAAATACCTTAAACCTTAATATCCGAATTAGAACTTCATCATACAAG
7710_donor          CTTATTTCTGCAGAAAATACCTTAAACCTTAATATCCGAATTAGAACTTCATCATACAAG
M32_outgroup        CTTATTTCTGCAGAAAATACCTTAAACCTTAATATCCGAATTAGAACTTCATCATACAAG
7710_outgroup       CTTATTTCTGCAGAAA-TACCTTAAACCTTAATATCCGAATTAGAACTTCATCATACAAG
                    **************** *******************************************

IAA_insertion       CATCAACATAACAAACTCCCACTGAAACTGTATATCCTTAATACCTTAACTGGCATGACA
M32_donor           CATCAACATAACAAACTCCCACTGAAACTGTATATCCTTAATACCTTAACTGGCATGACA
7710_donor          CATCAACATAACAAACTCCCACTGAAACTGTATATCCTTAATACCTTAACTGGCATGACA
M32_outgroup        CATCAACATAACAAACTCCCACTGAAACTGTATATCCTTAATACCTTAACTGGCATGACA
7710_outgroup       CATCAACATAACAAACTCCCACTGAAACTGTATATCCTTAATACCTTAACTGGCATGACA
                    ************************************************************

IAA_insertion       CACATGAATAATGTGCTCATGAAATACTTTAAGTATTTAACTTTAGTAAGCGGATCCGCA
M32_donor           CACATGAATAATGTGCTCATGAAATACTTTAAGTATTTAACTTTAGTAAGCGGATCCGCA
7710_donor          CACATGAATAATGTGCTCATGAAATACTTTAAGTATTTAACTTTAGTAAGCGGATCCGCA
M32_outgroup        CACATGAATAATGTGCTCATGAAATACTTTAAGTATTTAACTTTAGTAAGCGGATCCGCA
7710_outgroup       CACATGAATAATGTGCTCATGAAATACTTTAAGTATTTAACTTTAGTAAGCGGATCCGCA
                    ************************************************************

IAA_insertion       ATAATGGAGTTGTATTATGATGCTTTAAGTACAATAATTTTACTCTTCAAACTTTCACAA
M32_donor           ATAATGGAGTTGTATTATGATGCTTTAAGTACAATAATTTTACTCTTCAAACTTTCACAA
7710_donor          ATAATGGAGTTGTATTATGATGCTTTAAGTACAATAATTTTACTCTTCAAACTTTCACAA
M32_outgroup        ATAATGGAGTTGTATTATGATGCTTTAAGTACAATAATTTTACTCTTCAAACTTTCACAA
7710_outgroup       ATAATGGAGTTGTATTATGATGCTTTAAGTACAATAATTTTACTCTTCAAACTTTCACA-
```

```
                    ***********************************************************

IAA_insertion       CACTAAAGCCATGATAAAAATTTAACCGTATTTGGATTTCAAGCTACCTTGACATCCACC
M32_donor           CACTAAAGCCATGATAAAAATTTAACCGTATTTGGATTTCAAGCTACCTTGACATCCACC
7710_donor          CACTAAAGCCATGATAAAAATTTAACCGTATTTGGATTTCAAGCTACCTTGACATCCACC
M32_outgroup        CACTAAAGCCATGATAAAAATTTAACCGTATTTGGATTTCAAGCTACCTTGACATCCACC
7710_outgroup       CACTAAAGCCATGATAAAAATTTAACCGTATTTGGATTTCAAGCTACCTTGACATCCACC
                    ************************************************************

IAA_insertion       AAAAACGGAGTCTAATAGCTGTAAGTTCTCTCTAATGGATCTCTTTAATGATACTTATTC
M32_donor           AAAAACGGAGTCTAATAGCTGTAAGTTCTCTCTAATGGATCTCTTTAATGATACTTATTC
7710_donor          AAAAACGGAGTCTAATAGCTGTAAGTTCTCTCTAATGGATCTCTTTAATGATACTTATTC
M32_outgroup        AAAAACGGAGTCTAATAGCTGTAAGTTCTCTCTAATGGATCTCTTTAATGATACTTATTC
7710_outgroup       AAAA-CGGAGTCTAATAGCTGTAAGTTCTCTCTAATG-ATCTCTTTAATGATACTTATTC
                    **** ******************************** **********************

IAA_insertion       TTGAGGTTGTTGAGTTTGCTCATTGGAATTTTCCTTTTGAGATCCTAAATAGGGGAATAA
M32_donor           TTGAGGTTGTTGAGTTTGCTCATTGGAATTTTCCTTTTGAGATCCTAAATAGGGGAATAA
7710_donor          TTGAGGTTGTTGAGTTTGCTCATTGGAATTTTCCTTTTGAGATCCTAAATAGGGGAATAA
M32_outgroup        TTGAGGTTGTTGAGTTTGCTCATTGGAATTTTCCTTTTGAGATCCTAAATAGGGGAATAA
7710_outgroup       TTGAGGTTGTTGAGTTTGCTCATTGGAATTTTCCTTTTGAGATCCTAAATAGGGGAATAA
                    ************************************************************

IAA_insertion       ATCCTTATCTTCCCCCAAACTCAACATCCTCAAAGGACAAGCATCTATTGTATCCAAAAA
M32_donor           ATCCTTATCTTCCCCCAAACTCAACATCCTCAAAGGACAAGCATCTATTGTATCCAAAAA
7710_donor          ATCCTTATCTTCCCCCAAACTCAACATCCTCAAAGGACAAGCATCTATTGTATCCAAAAA
M32_outgroup        ATCCTTATCTTCCCCCAAACTCAACATCCTCAAAGGACAAGCATCTATTGTATCCAAAAA
7710_outgroup       ATCCTTATCTTCCCCCAAACTCAACATCCTCAAAGGACAAGCATCTATTGTATCCAAAAA
                    ************************************************************

IAA_insertion       TTGATCCAATAGTGAGACTATAAAAAAAAATGTATCCTTTATTATTATCTTCATCAATAT
M32_donor           TTGATCCAATAGTGAGACTATAAAAAAAA-TGTATCCTTTATTATTATCTTCATCAATAT
7710_donor          TTGATCCAATAGTGAGACTATAAAAAAAA-TGTATCCTTTATTATTATCTTCATCAATAT
M32_outgroup        TTGATCCAATAGTGAGACTATAAAAAAAA-TGTATCCTTTATCATTATCTTCATCAATAT
7710_outgroup       TTGATCCAATAGTGAGACTATAAAAAAAA-TGTATCCTTTATCATTATCTTCATCAATAT
                    ***************************** ************ *****************

IAA_insertion       CTTTAGTGAAAGATGTCAAGTGAGCACTTTATATCTTTCATTGTTTCAATCTCTCTTCCT
M32_donor           CTTTAGTGAAAGATGTCAAGTGAGCACTTTATATCTTTCATTGTTTCAATCTCTCTTCCT
7710_donor          CTTTAGTGAAAGATGTCAAGTGAGCACTTTATATCTTTCATTGTTTCAATCTCTCTTCCT
M32_outgroup        CTTTAGTGAAAGATGTCAAGTGAGCACTTTATATCTTTCATTGTTTCAATCTCTCTTCCT
7710_outgroup       CTTTAGTGAAAGATGTCAAGTGAGCACTTTATATCTTTCATTGTTTCAATCTCTCTTCCT
                    ************************************************************

IAA_insertion       CCGAATCAACATGAGTAGTGAACTCATTTAGAGGTTCACTTCTCCATAACAACAATTTAG
M32_donor           CCGAATCAACATGAGTAGTGAACTCATTTAGAGGTTCACTTCTCCATAACAACAATTTAG
7710_donor          CCGAATCAACATGAGTAGTGAACTCATTTAGAGGTTCACTTCTCCATAACAACAATTTAG
M32_outgroup        CCGAATCAACATGAGTAGTGAACTCATTTAGAGGTTCACTTCTCCATAACAACAATTTAG
7710_outgroup       CCGAATCAACATGAGTAGTGAACTCATTTAGAG-TTCACTTCTCCATAACAA---TTTAG
                    ********************************* ******************   *****

IAA_insertion       GAAGAGATTTAAAAACAAATGCACAAGCAAGTCGTTAGACTCCTCCAATTTAAGTGCCCT
M32_donor           GAAGAGATTTAAAAACAAATGCACAAGCAAGTCGTTAGACTCCTCCAATTTAAGTGCCCT
7710_donor          GAAGAGATTTAAAAACAAATGCACAAGCAAGTCGTTAGACTCCTCCAATTTAAGTGCCCT
M32_outgroup        GAAGAGATTTAAAAACAAATGCACAAGCAAGTCGTTAGACTCCTCCAATTTAAGTGCCCT
7710_outgroup       GAAGAGATTTAAAA-CAAATGCACAAGCAAGTCGTTAGACTCCTCCAATTTAAGTGCCCT
                    ************** *********************************************

IAA_insertion       TAAATCATGAAACAATATCACAAATTTCCATAATGTGCTCCCTTAATTTATTTATTTTCC
M32_donor           TAAATCATGAAACAATATCACAAATTTCCATAATGTGCTCCCTTAATTTATTTATTTTCC
7710_donor          TAAATCATGAAACAATATCACAAATTTCCATAATGTGCTCCCTTAATTTATTTATTTTCC
M32_outgroup        TAAATCATGAAACAATATCACAAATTTCCATAATGTGCTCCCTTAATTTATTTATTTTCC
7710_outgroup       TAA-TCATGAAACAATATCACAAATT-CCATAATGTGCTCCCTTAATTTATTTATTTTCC
```

```
                    *** ********************** *********************************

IAA_insertion       CCTTAATACCTTTTTCCCTTATGGAATTTAAGTCCAACAAGGTTGTACTCAATTCCCCTG
M32_donor           CCTTAATACCTTTTTCCCTTATGGAATTTAAGTCCAACAAGGTTGTACTCAATTCCCCTG
7710_donor          CCTTAATACCTTTTTCCCTTATGGAATTTAAGTCCAACAAGGTTGTACTCAATTCCCCTG
M32_outgroup        CCTTAATACCTTTTTCCCTTATGGAATTTAAGTCCAACAAGGTTGTACTCAATTCCCCTA
7710_outgroup       CCTTAATACCTTTTTCCCTTATGGAATTTAAGTCCAACAAGGTTGTACTCA-TTCCCCTA
                    *************************************************** ******* 

IAA_insertion       ATCTTTATAGCAAAACACTTTAAGCTCTTTAGAGAACAATTTGACATCTTTCTCTTTCAT
M32_donor           ATCTTTATAGCAAAACACTTTAAGCTCTTTAGAGAACAATTTGACATCTTTCTCTTTCAT
7710_donor          ATCTTTATAGCAAAACACTTTAAGCTCTTTAGAGAACAATTTGACATCTTTCTCTTTCAT
M32_outgroup        ATCTTTATAGCAAAACACTTTAAGCTCTTTAGAGAACAATTTGACATCTTTCTCTTTCAT
7710_outgroup       ATCTTTATAGCAAA-CACTTTAAGCTCTTTAGAGAACAATTTGACATCTTTCTCTTTCAT
                    ************** *********************************************

IAA_insertion       TTGGACATACATAATGTCCCCTAATAACTCCGAAATTGTACTTTAAATTGCACTTTACAT
M32_donor           TTGGACATACATAATGTCCCCTAATAACTCCGAAATTGTACTTTAAATTGCACTTTACAT
7710_donor          TTGGACATACATAATGTCCCCTAATAACTCCGAAATTGTACTTTAAATTGCACTTTACAT
M32_outgroup        TTGGACATACATAATGTCCCCTAATAACTCCGAAATTGTACTTTAAATTGCACTTTACAT
7710_outgroup       TTGGACATACATAATGTCCCCTAATAACTCCGAAATTGTACTTTAAATTGCACTTTACAT
                    ************************************************************

IAA_insertion       GATCATATACGACCTATCCAATTTGAGCATTCTCACTTCTCCATATTCCCCTTTAGTAGG
M32_donor           GATCATATACGACCTATCCAATTTGAGCATTCTCACTTCTCCATATTCCCCTTTAGTAGG
7710_donor          GATCATATACGACCTATCCAATTTGAGCATTCTCACTTCTCCATATTCCCCTTTAGTAGG
M32_outgroup        GATCATATACGACCTATGCAATTTGAGCATTCTCACTTCTCCATATTCCCCTTTAGTAGG
7710_outgroup       GATCATATACG-CCTATGCAATTTGAGCATTCTC-TTCTCCATATTCCCCTTTAGTAGG
                    *********** ***** **************** *************************

IAA_insertion       AGCACTAAAATCCATAAGAGTTGCGAGTGCTCAACTCGAAAAGCAAGGTCTGATCTAGAT
M32_donor           AGCACTAAAATCCATAAGAGTTGCGAGTGCTCAACTCGAAAAGCAAGGTCTGATCTAGAT
7710_donor          AGCACTAAAATCCATAAGAGTTGCGAGTGCTCAACTCGAAAAGCAAGGTCTGATCTGGAT
M32_outgroup        AGCACTAAAATCCGTAAGAGTTGCGAGTGCTCAACTCGAAAAGCAAGGTCTGATCTAGAT
7710_outgroup       --CACTAAAATCCGTAAGAGTTGCGAGTGCTCAACTCGAAAAGCAAGGTCTGATCTAGAT
                      *********** ****************************************** ***

IAA_insertion       CCATGCAAAAAAAAAAAAAAAA-CCAAAACCTTCAATTTATTATCCTTAATCAGATTAAT
M32_donor           CCATGCAAAAAAAAAAAAAAA--CCAAAACCTTCAATTTATTATCCTTAATCAGATTAAT
7710_donor          CCATGCAAAAAAAAAAAAAAAAACCAAAACCTTCAATTTATTATCCTTAATCAGATTAAT
M32_outgroup        CCATGCAAAAAAAAAAAAAA---CCAAAACCTTCAATTTATTATCCTTAATCAGATTAAT
7710_outgroup       CCATGCAAAAAAAAAAAA-----CCAAAACCTTCA-TTTATTATCCTTAATCAGATTAAT
                    ******************     ************ ************************

IAA_insertion       ATATTGACAATGTATCCCATCATGTACAAGAGACCTAACACAACATTAATTTTTT-AGTC
M32_donor           ATATTGACAATGTATCCCATCATGTACAAGAGACCTAACACAACATTAATTTTTT-AGTC
7710_donor          ATATTGACAATGTATCCCATCATGTACAAGAGACCTAACACAACATTAATTTTTT-AGTC
M32_outgroup        ATATTGACAATGTATCCCATCATGTACAAGAGACCTAACACAACATTAATTTTTTTAGTC
7710_outgroup       ATATTGACAATGTATCCCATCATGTACAAGAGACCTAACACAACATTAATTTTTTTAGTC
                    ******************************************************* ****

IAA_insertion       TTTGGACAGAAAAATTAACTTGTAAGTGGTCCTCTTGGTGTAGTAATCAAATACTAACAA
M32_donor           TTTGGACAGAAAAATTAACTTGTAAGTGGTCCTCTTGGTGTAGTAATCAAATACTAACAA
7710_donor          TTTGGACAGAAAAATTAACTTGTAAGTGGTCCTCTTGGTGTAGTAATCAAATACTAACAA
M32_outgroup        TTTGGACAGACAAATTAACTTGTAAGTGGTCCTCTTGGTGTAGTAATCAAATACTAACAA
7710_outgroup       TTTGGACAGACAAATTAACTTGTAAGTGGTCCTCTTGGTGTAGTAATCAAATACTAACAA
                    ********** *************************************************

IAA_insertion       TAAATTCATGTCAATGTTAAGCCTTTCTTTGGACCGACTTAACAACGCATGAAAAACCGA
M32_donor           TAAATTCATGTCAATGTTAAGCCTTTCTTTGGACCGACTTAACAACGCATGAAAAACCGA
7710_donor          TAAATTCATGTCAATGTTAAGCCTTTCTTTGGACCGACTTAACAACGCATGAAAAACCGA
M32_outgroup        TAAATTCATGTCAATGTTAAGCCTTTCTTTGGATCGACTTAACAACGCATGAAAAACCGA
7710_outgroup       TAAATTCATGTCAATGTTAAGCCTTTCTTTGGATCGACTTAAC---GCATGAAAAACCGA
```

```
                    ********************************* *********   **************

IAA_insertion       ACAATTGTTACATATTTATTACCACAAGCATGTTTAATTTCCGGCCAAATAATAGCCTTC
M32_donor           ACAATTGTTACATATTTATTACCACAAGCATGTTTAATTTCCGGCCAAATAATAGCCTTC
7710_donor          ACAATTGTTACATATTTATTACCACAAGCATGTTTAATTTCCGGCCAAATAATAGCCTTC
M32_outgroup        ACAATTGTTACATATTTATTACCACAAGCATGTTTAATTTCCGGCCAAATAATAGCCTTC
7710_outgroup       ACAATTGTTACATATTTATTACCACAAGCATGTTTAATTTCCGGCCAAATAATAGCCTTC
                    ************************************************************

IAA_insertion       CTTTGGGCCGACCATTATCCGCATGAAAACTAAACACACACAGGTATCTCAATATTTCAA
M32_donor           CTTTGGGCCGACCATTATCCGCATGAAAACTAAACACACACAGGTATCTCAATATTTCAA
7710_donor          CTTTGGGCCGACCATTATCCGCATGAAAACTAAACACACACAGGTATCTCAATATTTCAA
M32_outgroup        CTTTGGGCCGACCATTATCCGCATGAAAACTAAACACACACAGGTATCTCAATATTTCAA
7710_outgroup       CTTTGGGCCGACCATTATCCGCATGAAAACTAAACACACACAGGTATCTCAATATTTCAA
                    ************************************************************

IAA_insertion       ATTAATTGATCTACACAAAAGAGGTTACTTTGGCAACATATTGTTTCAATCAATTAATCC
M32_donor           ATTAATTGATCTACACAAAAGAGGTTACTTTGGCAACATATTGTTTCAATCAATTAATCC
7710_donor          ATTAATTGATCTACACAAAAGAGGTTACTTTGGCAACATATTGTTTCAATCAATTAATCC
M32_outgroup        ATTAATTGATCTACACAAAAGAGGTTACTTTGGCAACATATTGTTTCAATCAATTAATCC
7710_outgroup       -TTAATTGATCTACACAAAAGAG--TACTTTGGCAACATAT-GTTTCAATCAATTAATCC
                     **********************  **************** ******************

IAA_insertion       AAAATACTAGACCCAACTTAATAACCAAATTGCCAATTATTACAATCCGAAATATATGAT
M32_donor           AAAATACTAGACCCAACTTAATAACCAAATTGCCAATTATTACAATCCGAAATATATGAT
7710_donor          AAAATACTAGACCCAACTTAATAACCAAATTGCCAATTATTACAATCCGAAATATATGAT
M32_outgroup        AAAATACTAGACCCAACTTAATAACCAAATTGCCAATTATTACAATCCGAAATATATGAT
7710_outgroup       AAAATACTAGACCCAACTTAATAACCAAATTGCCAAT-ATTACAATCCGAA-TATATGA-
                    ************************************* ************* *******

IAA_insertion       GAATCCATAATTAGTCATGTGTTTATGAATCCATGACCTCAAATAGATAAGATGAAGCGC
M32_donor           GAATCCATAATTAGTCATGTGTTTATGAATCCATGACCTCAAATAGATAAGATGAAGCGC
7710_donor          GAATCCATAATTAGTCATGTGTTTATGAATCCATGACCTCAAATAGATAAGATGAAGCGC
M32_outgroup        GAATCCATAATTAGTCATGTGTTTATGAATCCATGACCTCAAATAGATAAGATGAAGCGC
7710_outgroup       --ATCCATAATTAGTCATGTGTTTATGAATCCATGACCTCAA-TAGATAAGATGAAGCGC
                      **************************************** *****************

IAA_insertion       AAAATAAGTTGTAAACTGTAACACTAAAAATTAATCGATGGATTCATAATTCATAGCAAC
M32_donor           AAAATAAGTTGTAAACTGTAACACTAAAAATTAATCGATGGATTCATAATTCATAGCAAC
7710_donor          AAAATAAGTTGTAAACTGTAACACTAAAAATTAATCGATGGATTCATAATTCATAGCAAC
M32_outgroup        AAAATAAGTTGTAAACTGTAACACTAAAAATTAATCGATGGATTCATAATTCATAGCAAC
7710_outgroup       AAA-TAAGTTGTAAACTGTAACACTAAAA-TTAATCGCTGGATTCATAATTCATAGCACA
                    *** ************************* ******* ********************

IAA_insertion       ACAGCAGCGACTAACAATTGAATTAAGCAATAAATTGTTCTGTCTTATATATGGAGTAAG
M32_donor           ACAGCAGCGACTAACAATTGAATTAAGCAATAAATTGTTCTGTCTTATATATGGAGTAAG
7710_donor          ACAGCAGCGACTAACAATTGAATTAAGCAATAAATTGTTCTGTCTTATATATGGAGTAAG
M32_outgroup        ACAGCAGCGACTAACAATTGAATTAAGCAATAAATTGTTCTGTCTTATATATGGAGTAAG
7710_outgroup       CCAGCA-CGACTAACAATTGAATTAAGCAATAAATT--TCTGTCTTATATATGGAGTAAG
                     ***** *****************************  **********************

IAA_insertion       ATTAAAATACCAAAGACTTTAACC--ACCAACACCCAGGCCAGGTCAAAAAATCAACCGA
M32_donor           ATTAAAATACCAAAGACTTTAACC--ACCAACACCCAGGCCAGGTCAAAAAATCAACCGA
7710_donor          ATTAAAATACCAAAGACTTTAACC--ACCAACACCCAGGCCAGGTCAAAAAATCAACCGA
M32_outgroup        ATTAAAATACCAAAGACTTTAACC--ACCAACACCCAGGCCAGGTCAAAAAATCAACCGA
7710_outgroup       ATTAAAA--CCAAAGACTTTAACCCCACCAACACCCAGGCCAGGTCAAAAAATCAACCGA
                    *******  ***************  **********************************

IAA_insertion       AATAAATAAAGTCAAGCTACATATATCTAGTGACTTTATCGATTCTAATCTTAAGAAAGG
M32_donor           AATAAATAAAGTCAAGCTACATATATCTAGTGACTTTATCGATTCTAATCTTAAGAAAGG
7710_donor          AATAAATAAAGTCAAGCTACATATATCTAGTGACTTTATCGATTCTAATCTTAAGAAAGG
M32_outgroup        AATAAATAAAGTCAAGCTACATATATCTAGTGACTTTATCGATTCTAATCTTAAGAAAGG
7710_outgroup       AATAAATAA-GTCAAGCTACATATATCTAGTGACTTTATCGATTCTAATCTTAAGAAAGG
```

```
                  ********* **************************************************

IAA_insertion     AATCACCAAATCAATGCCAAAACTAAAAAATAATCAGAACGATGACCAACCATATTTGAA
M32_donor         AATCACCAAATCAATGCCAAAACTAAAAAATAATCAGAACGATGACCAACCATATTTGAA
7710_donor        AATCACCAAATCAATGCCAAAACTAAAAAATAATCAGAACGATGACCAACCATATTTGAA
M32_outgroup      AATCACCAAATCAATGCCAAAACTAAAAAATAATCAGAACGATGACCAACCATATTTGAA
7710_outgroup     AATCACCAAATCAATGCCAAAACTAAAAAATAATCAGAACGATGACCA-CCATATTTGAA
                  ************************************************ ***********

IAA_insertion     GCACCCACAAAGCCTCATATGTCGAAATAAAATATCTGAAGTAAACTGTAAGGGATATTC
M32_donor         GCACCCACAAAGCCTCATATGTCGAAATAAAATATCTGAAGTAAACTGTAAGGGATATTC
7710_donor        GCACCCACAAAGCCTCATATGTCGAAATAAAATATCTGAAGTAAACTGTAAGGGATATTC
M32_outgroup      GCACCCACAAAGCCTCATATGTCGAAATAAAATATCTGAAGTAAACTGTAAGGGATATTC
7710_outgroup     GCACCCACAAAGCCTCATATGTCGAAATAAAATATCTGAAGTAAACTGTAAGGGATATTC
                  ************************************************************

IAA_insertion     AATTAAAGCATAATAGGGAATCAAAACCCAAATTAAATTTGGATGCTAACAAAGCCAAAA
M32_donor         AATTAAAGCATAATAGGGAATCAAAACCCAAATTAAATTTGGATGCTAACAAAGCCAAAA
7710_donor        AATTAAAGCATAATAGGGAATCAAAACCCAAATTAAATTTGGATGCTAACAAAGCCAAAA
M32_outgroup      AATTAAAGCATAATAGGGAATCAAAACCCAAATTAAATTTGGATGCTAACAAAGCCAAAA
7710_outgroup     AATTAA-GCATAATAGGGAATCAAAACCCAAATTAAATTTGGATGCTAACAAAGCCAAAA
                  ****** *****************************************************

IAA_insertion     ACATACACACACAACTGTCAACTGGATCGAATTGTGCATGTCAAATTGTCAATCATGACA
M32_donor         ACATACACACACAACTGTCAACTGGATCGAATTGTGCATGTCAAATTGTCAATCATGACA
7710_donor        ACATACACACACAACTGTCAACTGGATCGAATTGTGCATGTCAAATTGTCAATCATGACA
M32_outgroup      ACATACACACACAACTGTCAACTGGATCGAATTGTGCATGTCAAATTGTCAATCATGACA
7710_outgroup     ACATACACACACAACTGTCAACTGGATCGA-TTGTGCATGTCAA-TTGTCAATCATGACA
                  ****************************** ************* ***************

IAA_insertion     ATTGAGAAATAATTGATAACATACATCTGCCTTTTAGGCCTCGTCAATTAACAAATACTC
M32_donor         ATTGAGAAATAATTGATAACATACATCTGCCTTTTAGGCCTCGTCAATTAACAAATACTC
7710_donor        ATTGAGAAATAATTGATAACATACATCTGCCTTTTAGGCCTCGTCAATTAACAAATACTC
M32_outgroup      ATTGAGAAATAATTGATAACATACATCTGCCTTTTAGGCCTCGTCAATTAACAAATACTC
7710_outgroup     ATTGAGAAATAATTGATAACATACATCTGCCTTTTAGGCCTCGTCAATAAC--AATACTC
                  ************************************************ *   *******

IAA_insertion     AAACTCAATAGTTGATCACAAAACAAATTCGAAGTTCTGAAAGAAAAATACTTCGTACCA
M32_donor         AAACTCAATAGTTGATCACAAAACAAATTCGAAGTTCTGAAAGAAAAATACTTCGTACCA
7710_donor        AAACTCAATAGTTGATCACAAAACAAATTCGAAGTTCTGAAAGAAAAATACTTCGTACCA
M32_outgroup      AAACTCAATAGTTGATCACAAAACAAATTCGAAGTTCTGAAAGAAAAATACTTCGTACCA
7710_outgroup     AAACTCAATAGTTGATCACAAAACAAATTCGA-GTTCTGAAAGAAAAATACTTCGTACCA
                  ******************************** ***************************

IAA_insertion     AATAATTTAATATGAAGAAGGAGATCCATAAAGAATTGGACTTGTTGGGATTAAACCCTA
M32_donor         AATAATTTAATATGAAGAAGGAGATCCATAAAGAATTGGACTTGTTGGGATTAAACCCTA
7710_donor        AATAATTTAATATGAAGAAGGAGATCCATAAAGAATTGGACTTGTTGGGATTAAACCCTA
M32_outgroup      AATAATTTAATATGAAGAAGGAGATCCATAAAGAATTGGACTAGTTGGGATTAAACCCTA
7710_outgroup     A-TAATTTAATATGAAGAAGGAGATCCATAAAGAATTGGACTAGTTGGGATTAA-CCCTA
                  * **************************************** *********** *****

IAA_insertion     ATTCATATCAAATTAGTCGGTAATAATAGATCTGCAGAAGCATACCTGAATCCATGACGA
M32_donor         ATTCATATCAAATTAGTCGGTAATAATAGATCTGCAGAAGCATACCTGAATCCATGACGA
7710_donor        ATTCATATCAAATTAGTCGGTAATAATAGATCTGCAGAAGCGTACCTGAATCCATGACGA
M32_outgroup      ATTCATATCAAATTAGTCGGTAATAATAGATCTGCAGAAGCATACCTGAATCCATGACGA
7710_outgroup     ATTCATATCAAATTAGTCG-TAATAATAGATCTGCAGAAGCATACCTGAATCCATGACGA
                  ******************* ********************* ******************

IAA_insertion     AGATCGGCGGTGGGTTTTGATCTTCCAATTCTTTATGGCTCCTTAGGGTTTCTACTGATG
M32_donor         AGATCGGCGGTGGGTTTTGATCTTCCAATTCTTTATGGCTCCTTAGGGTTTCTACTGATG
7710_donor        AGATCGGCGGTGGGTTTTGATCTTCCAATTCTTTATGGCTCCTTAGGGTTTCTACTGATG
M32_outgroup      AGATCGGCGGTGGGTTTTGATCTTCCAATTCTTTATGGCTCCTTAGGGTTTCTACTGATG
7710_outgroup     AGATCGGCGG--GGTTTTGATCTTCCAATTCTTTATGGCTCCTTAGG-TTTCTACTGATG
```

```
                   **********  *********************************** ************

IAA_insertion      ATGGGATGTCAGTGAGAATAGGAAAACCATATGAATCGGGGACCATAACCCTTAATGTCT
M32_donor          ATGGGATGTCAGTGAGAATAGGAAAACCATATGAATCGGGGACCATAACCCTTAATGTCT
7710_donor         ATGGGATGTCAGTGAGAATAGGAAAACCATATGAATCGGGGACCATAACCCTTAATGTCT
M32_outgroup       ATGGGATGTCAGTGAGAATAGGAAAACCATATAAATCGGGGACCATAACCCTTAATGTCT
7710_outgroup      --GGGATGTCAGTGAGAATAGGAAAACCATATAAATCGGGGACCATAACCCTTAATGTCT
                     ****************************** ***************************

IAA_insertion      ATTTATATACATAGACTCCTTCCTAATCCGCCCATCGTGAATAAGGAAAGGCCCATCGGT
M32_donor          ATTTATATACATAGACTCCTTCCTAATCCGCCCATCGTGAATAAGGAAAGGCCCATCGGT
7710_donor         ATTTATATACATAGACTCCTTCCTAATCCGCCCATCGTGAATAAGGAAAGGCCCATCGGT
M32_outgroup       ATTTATATACATAGACTCCTTCCTAATCCGCCCATCGTGAATAAGGAAAGGCCCATCGGT
7710_outgroup      ATTTATATACATAGACTCCTTCCTAATCC-CCCATCGTGAATAAGGAAAGGCCCATCG-T
                   ***************************** **************************** *

IAA_insertion      ATCTACACAAATAAAAAACTGATCCCACACTCTATAAAAGACGTAAATAGGCCCAATAAT
M32_donor          ATCTACACAAATAAAAAACTGATCCCACACTCTATAAAAGACGTAAATAGGCCCAATAAT
7710_donor         ATCTACACAAATAAAAAACTGATCCCACACTCTATAAAAGACGTAAATAGGCCCAATAAT
M32_outgroup       ATCTACACAAATAAAAAACTGATCCCACACTCTATAAAAGACGTAAATAGGCCCAATAAT
7710_outgroup      ATCTACACAAATAAAAAACTGATCCCACACTCTATAAAA--CGTAAATAG-CCCAATAAT
                   ***************************************  ********* *********

IAA_insertion      AATTACTTAATTGGATCACTTTAGTTTTGGGCCACACCGTATGATAGCACATAATACAAT
M32_donor          AATTACTTAATTGGATCACTTTAGTTTTGGGCCACACCGTATGATAGCACATAATACAAT
7710_donor         AATTACTTAATTGGATCACTTTAGTTTTGGGCCACACCGTATGATAGCACATAATACAAT
M32_outgroup       AATTACTTAATTGGATCACTTTAGTTTTGGGCCACACCGTATGATAGCACATAATACAAT
7710_outgroup      AATTACTTAATTGGATCACTTTAGTTTTGGGCCACACCGTATGATAGCACATAATACAAT
                   ************************************************************

IAA_insertion      TATAACTGAATTGCACACGTATTTGTATTTAGGTCCATAAAATTTCCAACATGGCC
M32_donor          TATAACTGAATTGCACACGTATTTGTATTTAGGTCCATAAAATTTCCAACA-----
7710_donor         TATAACTGAATTGCACACGTATTTGTATTTAGGTCCATAAAATTTCCAACA-----
M32_outgroup       TATAACTGAATTGCACACGTATTTGTATTTAGGTCCATAAAATTTCCAACA-----
7710_outgroup      TATAACTGAATTGCACACGTATT-GTATTTAGGTCCATAAAATTTCCAACA-----
                   *********************** ***************************
```

Figure S2. Maximum Likelihood tree showing relatedness of retrotransposons similar to the one inserted in *IAA16*. Names correspond to the descriptions in the caption of Table S5.

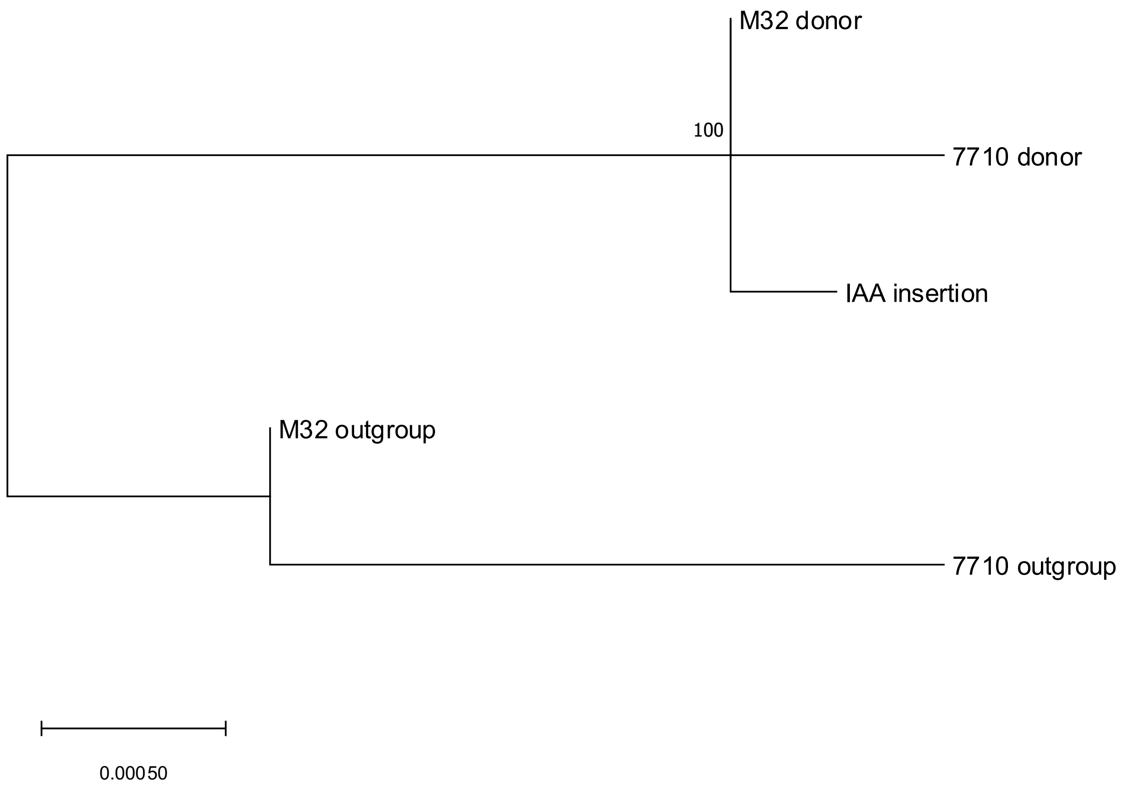

Figure S3. *Arabidopsis thaliana* plants (genotype Col 0) plants expressing *BsIAA16* alleles (*BsIAA16*$_{WT}$, 7710 2-6-2; *BsIAA16*$_{MUT}$, M32 2-3-5) or with no transgene (02A) either untreated or treated with 140 g dicamba ha$^{-1}$. Photo taken 14 days after treatment.

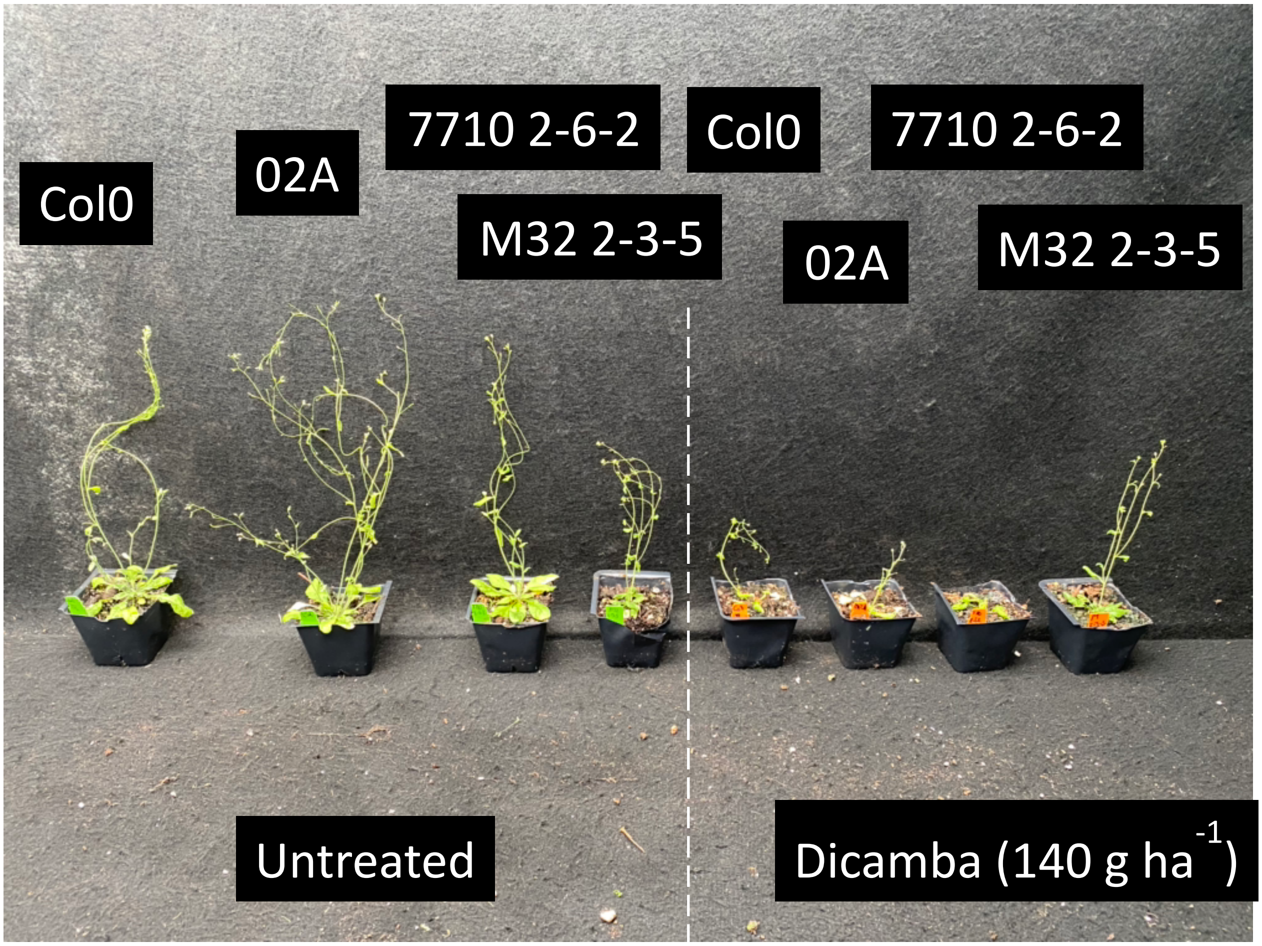

Table S6. KASP primer set for identification of M32 allele of IAA16. The primer that binds to the wildtype has a FAM tail (green) and the primer that binds the M32 allele has a HEX tail (orange). Thermocycler conditions are 94C for 15 min; 10 cycles of 94C for 20 seconds, 61 C for 60 seconds and decreasing 0.6 C each cycle; 40 cycles of 94C for 20 seconds, 55 C for 60 seconds, quantifying FAM and HEX fluorescence at each round at 30C.

| Primer name | Sequence (5'-3') |
|---|---|
| BsIAA16_intron1F_WT_KASP | GAAGGTGACCAAGTTCATGCTCTTCAGGACACAAGTTGTAGGT |
| BsIAA16_intron1F_M32_KASP | GAAGGTCGGAGTCAACGGATTGCCACACCGTATGATAGCAC |
| BsIAA16_484R_KASP | CGCTTGTGATGGCATTGCTA |

Table S7. Polymerase chain reaction primers used to amplify *BsIAA16* from cDNA libraries. Thermocycler conditions were 95 C for 5 minutes; 35 cycles of 95 C for 15 seconds, 60 C for 15 seconds, 72 C for 1 minute; 72 C for 5 minutes.

| Primer name | Sequence (5'-3') |
|---|---|
| BsIAA16_1F | ATGTTGAGTAACGAGAGAGAC |
| BsIAA16_843R | TCAGCTTCTGTTCTTGCACT |

Table S8. Polymerase chain reaction primers used to clone *BsAUX/IAA16* into pFGC5941. Thermocycler conditions were 95 C for 5 minutes; 35 cycles of 95 C for 15 seconds, 60 C for 15 seconds, 72 C for 1 minute; 72 C for 5 minutes. Tails used for In-Fusion cloning are colored in green.

| Primer name | Sequence (5'-3') |
|---|---|
| BsIAA16_clone_1F | TTACCATGGGGCGCGATGTTGAGTAACGAGAGAGACAAG |
| BsIAA16_clone_843R | GACTCACCTAGGATCTCAGCTTCTGTTCTTGCACTTC |

Table S9. Polymerase chain reaction primers used to amplify a section of pFGC5941 that contains the transgene of interest. Thermocycler conditions were 95 C for 5 minutes; 35 cycles of 95 C for 15 seconds, 60 C for 15 seconds, 72 C for 2 minutes; 72 C for 5 minutes. Positive samples should show an amplicon of ~400bp + the length of your transgene.

| Primer name | Sequence (5'-3') |
|---|---|
| pFGC_F | CCAACCACGTCTTCAAAGCA |
| pFGC_R | GGCGTCTCGCATATCTCATT |

Table S10. Quantitative PCR primers to quantify expression of various *Arabidopsis thaliana* and *Bassia scoparia* genes. Thermocycler conditions for all primer sets are 95C for 3 minutes; 40 cycles of 95C for 15 seconds, 60C for 30 seconds, quantifying SYBR fluorescence after each round at 60C.

| Primer name | Sequence (5'-3') |
|---|---|
| BsIAA16_qPCR_393R | GCGACGATGTTCTTCCTGAA |
| BsIAA16_qPCR_209F | GCAACCAAAACTACAGCGGA |
| BsSAUR21_qPCR_1F | ATGGCAATCCGATTTCCTTCAG |
| BsSAUR21_qPCR_84R | GATGTACCCTTTTGGAACTGCTT |
| BsActin_R | ATGAGAGAACGGCCTGAATG |
| BsActin_F | GAGCATCCTGTCTTACTGACTG |
| AtCyclophilin_qPCR_R | AATCGGCAACAACCACAGGC |
| AtCyclophilin_qPCR_F | GTCTGATAGAGATCTCACGT |
| AtGH3.3_qPCR_R | CGTCATTTGGAGGATTGGTTTG |
| AtGH3.3_qPCR_F | GAGAGCAAGGAACTCGTGTTAT |
| AtIAA19_qPCR_R | CTCCGTGAAAGCTCTCTTCTTC |
| AtIAA19_qPCR_F | AGGACTCGGGCTTGAGATAA |

Table S11. Phenotypic response of kochia from a $F_3$ populations segregating for herbicide resistance. Visual injury was rated 21 days after treatment on a scale from 0-100 and survival was noted (A=alive, D=dead).

| **Sample** | **Injury percentage** | **Survival** |
|---|---:|---|
| 7710xM32-4-1-1-1 | 50 | A |
| 7710xM32-4-1-1-2 | 0 | A |
| 7710xM32-4-1-1-3 | 0 | A |
| 7710xM32-4-1-1-4 | 100 | D |
| 7710xM32-4-1-1-5 | 0 | A |
| 7710xM32-4-1-1-6 | 100 | D |
| 7710xM32-4-1-1-7 | 0 | A |
| 7710xM32-4-1-1-8 | 0 | A |
| 7710xM32-4-1-1-9 | 0 | A |
| 7710xM32-4-1-1-10 | 0 | A |
| 7710xM32-4-1-1-11 | 0 | A |
| 7710xM32-4-1-1-12 | 0 | A |
| 7710xM32-4-1-1-13 | 0 | A |
| 7710xM32-4-1-1-14 | 0 | A |
| 7710xM32-4-1-1-15 | 100 | D |
| 7710xM32-4-1-1-16 | 100 | D |

| | | |
|---|---|---|
| 7710xM32-4-1-1-17 | 100 | D |
| 7710xM32-4-1-1-18 | 20 | A |
| 7710xM32-4-1-1-19 | 0 | A |
| 7710xM32-4-1-1-20 | 100 | D |
| 7710xM32-4-1-1-21 | 100 | D |
| 7710xM32-4-1-1-22 | 100 | D |
| 7710xM32-4-1-1-23 | 0 | A |
| 7710xM32-4-1-1-24 | 30 | A |
| 7710xM32-4-1-1-25 | 0 | A |
| 7710xM32-4-1-1-26 | 0 | A |
| 7710xM32-4-1-1-27 | 50 | A |
| 7710xM32-4-1-1-28 | 0 | A |
| 7710xM32-4-1-1-29 | 0 | A |
| 7710xM32-4-1-1-30 | 0 | A |
| 7710xM32-4-1-1-31 | 0 | A |
| 7710xM32-4-1-1-32 | 0 | A |
| 7710xM32-4-1-1-33 | 0 | A |
| 7710xM32-4-1-1-34 | 30 | A |
| 7710xM32-4-1-1-35 | 30 | A |
| 7710xM32-4-1-1-36 | 0 | A |
| 7710xM32-4-1-1-37 | 0 | A |
| 7710xM32-4-1-1-38 | 0 | A |
| 7710xM32-4-1-1-39 | 0 | A |
| 7710xM32-4-1-1-40 | 0 | A |
| 7710xM32-4-1-1-41 | 0 | A |
| 7710xM32-4-1-1-42 | 100 | D |
| 7710xM32-4-1-1-43 | 0 | A |
| 7710xM32-4-1-1-44 | 0 | A |
| 7710xM32-4-1-1-45 | 100 | D |
| 7710xM32-4-1-1-46 | 20 | A |
| 7710xM32-4-1-1-47 | 30 | A |
| 7710xM32-4-1-1-48 | 0 | A |
| 7710xM32-4-1-1-49 | 100 | D |
| 7710xM32-4-1-1-50 | 100 | D |
| 7710xM32-4-1-1-51 | 100 | D |
| 7710xM32-4-1-1-52 | 100 | D |
| 7710xM32-4-1-1-53 | 50 | A |
| 7710xM32-4-1-1-54 | 40 | A |
| 7710xM32-4-1-1-55 | 30 | A |

| | | |
|---|---|---|
| 7710xM32-4-1-1-56 | 20 | A |
| 7710xM32-4-1-1-57 | 50 | A |
| 7710xM32-4-1-1-58 | 20 | A |
| 7710xM32-4-1-1-59 | 40 | A |
| 7710xM32-4-1-1-60 | 30 | A |
| 7710xM32-4-1-1-61 | 40 | A |
| 7710xM32-4-1-1-62 | 50 | A |
| 7710xM32-4-1-1-63 | 40 | A |
| 7710xM32-4-1-1-64 | 50 | A |
| 7710xM32-4-1-1-65 | 30 | A |
| 7710xM32-4-1-1-66 | 0 | A |
| 7710xM32-4-1-1-67 | 50 | A |
| 7710xM32-4-1-1-68 | 0 | A |
| 7710xM32-4-1-1-69 | 0 | A |
| 7710xM32-4-1-1-70 | 0 | A |
| 7710xM32-4-1-1-71 | 50 | A |
| 7710xM32-4-1-1-72 | 50 | A |
| 7710xM32-4-1-1-73 | 0 | A |
| 7710xM32-4-1-1-74 | 0 | A |
| 7710xM32-4-1-1-75 | 0 | A |
| 7710xM32-4-1-1-76 | 50 | A |
| 7710xM32-4-1-1-77 | 100 | D |
| 7710xM32-4-1-1-78 | 0 | A |
| 7710xM32-4-1-1-79 | 30 | A |
| 7710xM32-4-1-1-80 | 0 | A |
| 7710xM32-4-1-1-81 | 100 | D |
| 7710xM32-4-1-1-82 | 100 | D |
| 7710xM32-4-1-1-83 | 50 | A |
| 7710xM32-4-1-1-84 | 30 | A |
| 7710xM32-4-1-1-85 | 100 | D |
| 7710xM32-4-1-1-86 | 100 | D |
| 7710xM32-4-1-1-87 | 50 | A |
| 7710xM32-4-1-1-88 | 100 | D |
| 7710xM32-4-1-1-89 | 100 | D |
| 7710xM32-4-1-1-90 | 100 | D |
| 7710xM32-4-1-1-91 | 100 | D |
| 7710xM32-4-1-1-92 | 100 | D |
| 7710xM32-4-1-1-93 | 100 | D |
| 7710xM32-4-1-1-94 | 100 | D |

| | | |
|---|---|---|
| 7710xM32-4-1-1-95 | 30 | A |
| 7710xM32-4-1-1-96 | 100 | D |
| 7710xM32-4-1-1-97 | 100 | D |
| 7710xM32-4-1-1-98 | 0 | A |
| 7710xM32-4-1-1-99 | 100 | D |
| 7710xM32-4-1-1-100 | 60 | A |
| 7710xM32-4-1-1-101 | 0 | A |
| 7710xM32-4-1-1-102 | 0 | A |
| 7710xM32-4-1-1-103 | 0 | A |
| 7710xM32-4-1-1-104 | 0 | A |
| 7710xM32-4-1-1-105 | 100 | D |
| 7710xM32-4-1-1-106 | 50 | A |
| 7710xM32-4-1-1-107 | 100 | D |
| 7710xM32-4-1-1-108 | 0 | A |
| 7710xM32-4-1-1-109 | 100 | D |
| 7710xM32-4-1-1-110 | 100 | D |
| 7710xM32-4-1-1-111 | 0 | A |
| 7710xM32-4-1-1-112 | 100 | D |
| 7710xM32-4-1-1-113 | 30 | A |
| 7710xM32-4-1-1-114 | 100 | D |
| 7710xM32-4-1-1-115 | 0 | A |
| 7710xM32-4-1-1-116 | | No data |
| 7710xM32-4-1-1-117 | 20 | A |
| 7710xM32-4-1-1-118 | 100 | D |
| 7710xM32-4-1-1-119 | 30 | A |
| 7710xM32-4-1-1-120 | 20 | A |
| 7710xM32-4-1-1-121 | 50 | A |
| 7710xM32-4-1-1-122 | 0 | A |
| 7710xM32-4-1-1-123 | 100 | D |
| 7710xM32-4-1-1-124 | 30 | A |
| 7710xM32-4-1-1-125 | 100 | D |
| 7710xM32-4-1-1-126 | 100 | D |
| 7710xM32-4-1-1-127 | 50 | A |
| 7710xM32-4-1-1-128 | 30 | A |
| 7710xM32-4-1-1-129 | 20 | A |
| 7710xM32-4-1-1-130 | 20 | A |
| 7710xM32-4-1-1-131 | 20 | A |
| 7710xM32-4-1-1-132 | 100 | D |
| 7710xM32-4-1-1-133 | 20 | A |

| | | |
|---|---|---|
| 7710xM32-4-1-1-134 | 20 | A |
| 7710xM32-4-1-1-135 | 20 | A |
| 7710xM32-4-1-1-136 | 20 | A |
| 7710xM32-4-1-1-137 | 100 | D |
| 7710xM32-4-1-1-138 | 20 | A |
| 7710xM32-4-1-1-139 | 100 | D |
| 7710xM32-4-1-1-140 | 100 | D |
| 7710xM32-4-1-1-141 | 20 | A |
| 7710xM32-4-1-1-142 | 30 | A |
| 7710xM32-4-1-1-143 | | No data |
| 7710xM32-4-1-1-144 | 100 | D |
| 7710xM32-4-1-1-145 | 20 | A |
| 7710xM32-4-1-1-146 | 20 | A |
| 7710xM32-4-1-1-147 | 100 | D |
| 7710xM32-4-1-1-148 | 20 | A |
| 7710xM32-4-1-1-149 | 30 | A |
| 7710xM32-4-1-1-150 | 100 | D |
| 7710xM32-4-1-1-151 | 100 | D |
| 7710xM32-4-1-1-152 | 20 | A |
| 7710xM32-4-1-1-153 | 100 | D |
| 7710xM32-4-1-1-154 | 100 | D |
| 7710xM32-4-1-1-155 | 100 | D |
| 7710xM32-4-1-1-156 | 20 | A |
| 7710xM32-4-1-1-157 | 100 | D |
| 7710xM32-4-1-1-158 | 100 | D |
| 7710xM32-4-1-1-159 | 20 | A |
| 7710xM32-4-1-1-160 | 20 | A |
| 7710xM32-4-1-1-161 | 50 | A |
| 7710xM32-4-1-1-162 | 50 | A |
| 7710xM32-4-1-1-163 | 0 | A |
| 7710xM32-4-1-1-164 | 50 | A |
| 7710xM32-4-1-1-165 | 50 | A |
| 7710xM32-4-1-1-166 | 50 | A |
| 7710xM32-4-1-1-167 | 100 | D |
| 7710xM32-4-1-1-168 | 50 | A |
| 7710xM32-4-1-1-169 | 50 | A |
| 7710xM32-4-1-1-170 | 100 | D |
| 7710xM32-4-1-1-171 | 100 | D |
| 7710xM32-4-1-1-172 | 50 | A |

| | | |
|---|---|---|
| 7710xM32-4-1-1-173 | 100 | D |
| 7710xM32-4-1-1-174 | 30 | A |
| 7710xM32-4-1-1-175 | 50 | A |
| 7710xM32-4-1-1-176 | 50 | A |
| 7710xM32-4-1-1-177 | 100 | D |
| 7710xM32-4-1-1-178 | 50 | A |
| 7710xM32-4-1-1-179 | 100 | D |
| 7710xM32-4-1-1-180 | 50 | A |
| 7710xM32-4-1-1-181 | 0 | A |
| 7710xM32-4-1-1-182 | 100 | D |
| 7710xM32-4-1-1-183 | 30 | A |
| 7710xM32-4-1-1-184 | 50 | A |
| 7710xM32-4-1-1-185 | 20 | A |
| 7710xM32-4-1-1-186 | 0 | A |
| 7710xM32-4-1-1-187 | | No data |
| 7710xM32-4-1-1-188 | 50 | A |
| 7710xM32-4-1-1-189 | 20 | A |
| 7710xM32-4-1-1-190 | 0 | A |
| 7710xM32-4-1-1-191 | 100 | D |
| 7710xM32-4-1-1-192 | 100 | D |
| 7710xM32-4-1-1-193 | 0 | A |
| 7710xM32-4-1-1-194 | 50 | A |
| 7710xM32-4-1-1-195 | 100 | D |
| 7710xM32-4-1-1-196 | 20 | A |
| 7710xM32-4-1-1-197 | 30 | A |
| 7710xM32-4-1-1-198 | 0 | A |
| 7710xM32-4-1-1-199 | 30 | A |
| 7710xM32-4-1-1-200 | 0 | A |
| 7710xM32-4-1-1-201 | 0 | A |
| 7710xM32-4-1-1-202 | 30 | A |
| 7710xM32-4-1-1-203 | 0 | A |
| 7710xM32-4-1-1-204 | 30 | A |
| 7710xM32-4-1-1-205 | 30 | A |
| 7710xM32-4-1-1-206 | 30 | A |
| 7710xM32-4-1-1-207 | 30 | A |
| 7710xM32-4-1-1-208 | 100 | D |
| 7710xM32-4-1-1-209 | 30 | A |
| 7710xM32-4-1-1-210 | 30 | A |
| 7710xM32-4-1-1-211 | 30 | A |

| | | |
|---|---|---|
| 7710xM32-4-1-1-212 | 30 | A |
| 7710xM32-4-1-1-213 | 0 | A |
| 7710xM32-4-1-1-214 | 30 | A |
| 7710xM32-4-1-1-215 | 0 | A |
| 7710xM32-4-1-1-216 | 30 | A |
| 7710xM32-4-1-1-217 | 30 | A |
| 7710xM32-4-1-1-218 | 0 | A |
| 7710xM32-4-1-1-219 | 0 | A |
| 7710xM32-4-1-1-220 | 30 | A |
| 7710xM32-4-1-1-221 | 30 | A |
| 7710xM32-4-1-1-222 | 30 | A |
| 7710xM32-4-1-1-223 | 30 | A |
| 7710xM32-4-1-1-224 | 30 | A |
| 7710xM32-4-1-1-225 | 100 | D |
| 7710xM32-4-1-1-226 | 100 | D |
| 7710xM32-4-1-1-227 | 50 | A |
| 7710xM32-4-1-1-228 | 30 | A |
| 7710xM32-4-1-1-229 | 100 | D |
| 7710xM32-4-1-1-230 | 100 | D |
| 7710xM32-4-1-1-231 | 0 | A |
| 7710xM32-4-1-1-232 | 100 | D |
| 7710xM32-4-1-1-233 | 100 | D |
| 7710xM32-4-1-1-234 | 0 | A |
| 7710xM32-4-1-1-235 | 30 | A |
| 7710xM32-4-1-1-236 | 100 | D |
| 7710xM32-4-1-1-237 | 0 | A |
| 7710xM32-4-1-1-238 | 0 | A |
| 7710xM32-4-1-1-239 | 30 | A |
| 7710xM32-4-1-1-240 | 30 | A |
| 7710xM32-4-1-1-241 | 0 | A |
| 7710xM32-4-1-1-242 | 100 | D |
| 7710xM32-4-1-1-243 | 100 | D |
| 7710xM32-4-1-1-244 | 20 | A |
| 7710xM32-4-1-1-245 | 100 | D |
| 7710xM32-4-1-1-246 | 0 | A |
| 7710xM32-4-1-1-247 | 0 | A |
| 7710xM32-4-1-1-248 | 20 | A |
| 7710xM32-4-1-1-249 | 0 | A |
| 7710xM32-4-1-1-250 | 0 | A |

| | | |
|---|---|---|
| 7710xM32-4-1-1-251 | 0 | A |
| 7710xM32-4-1-1-252 | 20 | A |
| 7710xM32-4-1-1-253 | 100 | D |
| 7710xM32-4-1-1-254 | 0 | A |
| 7710xM32-4-1-1-255 | 20 | A |
| 7710xM32-4-1-1-256 | 100 | D |
| 7710xM32-4-1-1-257 | 0 | A |
| 7710xM32-4-1-1-258 | 0 | A |
| 7710xM32-4-1-1-259 | 20 | A |
| 7710xM32-4-1-1-260 | 100 | D |
| 7710xM32-4-1-1-261 | 40 | A |
| 7710xM32-4-1-1-262 | 30 | A |
| 7710xM32-4-1-1-263 | 100 | D |
| 7710xM32-4-1-1-264 | 40 | A |
| 7710xM32-4-1-1-265 | 100 | D |
| 7710xM32-4-1-1-266 | 30 | A |
| 7710xM32-4-1-1-267 | 100 | D |
| 7710xM32-4-1-1-268 | 20 | A |
| 7710xM32-4-1-1-269 | 100 | D |
| 7710xM32-4-1-1-270 | 30 | A |
| 7710xM32-4-1-1-271 | 30 | A |
| 7710xM32-4-1-1-272 | 100 | D |
| 7710xM32-4-1-1-273 | 40 | A |
| 7710xM32-4-1-1-274 | 100 | D |
| 7710xM32-4-1-1-275 | 30 | A |
| 7710xM32-4-1-1-276 | 20 | A |
| 7710xM32-4-1-1-277 | 30 | A |
| 7710xM32-4-1-1-278 | 30 | A |
| 7710xM32-4-1-1-279 | 100 | D |
| 7710xM32-4-1-1-280 | 100 | D |
| 7710xM32-4-1-1-281 | 30 | A |
| 7710xM32-4-1-1-282 | 30 | A |
| 7710xM32-4-1-1-283 | 30 | A |
| 7710xM32-4-1-1-284 | 30 | A |
| 7710xM32-4-1-1-285 | 100 | D |
| 7710xM32-4-1-1-286 | 30 | A |
| 7710xM32-4-1-1-287 | 30 | A |
| 7710xM32-4-1-1-288 | 0 | A |
| 7710-M32-4-5-10-1 | 100 | D |

| | | |
|---|---|---|
| 7710-M32-4-5-10-2 | 100 | D |
| 7710-M32-4-5-10-3 | 70 | A |
| 7710-M32-4-5-10-4 | 100 | D |
| 7710-M32-4-5-10-5 | 30 | A |
| 7710-M32-4-5-10-6 | 70 | A |
| 7710-M32-4-5-10-7 | | No data |
| 7710-M32-4-5-10-8 | 100 | D |
| 7710-M32-4-5-10-9 | 30 | A |
| 7710-M32-4-5-10-10 | 100 | D |
| 7710-M32-4-5-10-11 | 20 | A |
| 7710-M32-4-5-10-12 | 70 | A |
| 7710-M32-4-5-10-13 | 100 | D |
| 7710-M32-4-5-10-14 | 20 | A |
| 7710-M32-4-5-10-15 | 20 | A |
| 7710-M32-4-5-10-16 | 20 | A |
| 7710-M32-4-5-10-17 | 100 | D |
| 7710-M32-4-5-10-18 | 20 | A |
| 7710-M32-4-5-10-19 | 80 | A |
| 7710-M32-4-5-10-20 | 100 | D |
| 7710-M32-4-5-10-21 | 100 | D |
| 7710-M32-4-5-10-22 | 20 | A |
| 7710-M32-4-5-10-23 | | No data |
| 7710-M32-4-5-10-24 | 20 | A |
| 7710-M32-4-5-10-25 | 20 | A |
| 7710-M32-4-5-10-26 | 70 | A |
| 7710-M32-4-5-10-27 | 30 | A |
| 7710-M32-4-5-10-28 | 20 | A |
| 7710-M32-4-5-10-29 | 20 | A |
| 7710-M32-4-5-10-30 | 20 | A |
| 7710-M32-4-5-10-31 | 20 | A |
| 7710-M32-4-5-10-32 | | No data |
| 7710-M32-4-5-10-33 | 100 | D |
| 7710-M32-4-5-10-34 | 20 | A |
| 7710-M32-4-5-10-35 | 100 | D |
| 7710-M32-4-5-10-36 | 100 | D |
| 7710-M32-4-5-10-37 | 100 | D |
| 7710-M32-4-5-10-38 | 80 | A |
| 7710-M32-4-5-10-39 | 100 | D |
| 7710-M32-4-5-10-40 | | No data |

| | | |
|---|---|---|
| 7710-M32-4-5-10-41 | 100 | D |
| 7710-M32-4-5-10-42 | 30 | A |
| 7710-M32-4-5-10-43 | 100 | D |
| 7710-M32-4-5-10-44 | | No data |
| 7710-M32-4-5-10-45 | 100 | D |
| 7710-M32-4-5-10-46 | 100 | D |
| 7710-M32-4-5-10-47 | 20 | A |
| 7710-M32-4-5-10-48 | 80 | A |
| 7710-M32-4-5-10-49 | 30 | A |
| 7710-M32-4-5-10-50 | 30 | A |
| 7710-M32-4-5-10-51 | 100 | D |
| 7710-M32-4-5-10-52 | 30 | A |
| 7710-M32-4-5-10-53 | 30 | A |
| 7710-M32-4-5-10-54 | 100 | D |
| 7710-M32-4-5-10-55 | 100 | D |
| 7710-M32-4-5-10-56 | 100 | D |
| 7710-M32-4-5-10-57 | | No data |
| 7710-M32-4-5-10-58 | 50 | A |
| 7710-M32-4-5-10-59 | 100 | D |
| 7710-M32-4-5-10-60 | 100 | D |
| 7710-M32-4-5-10-61 | 80 | A |
| 7710-M32-4-5-10-62 | 80 | A |
| 7710-M32-4-5-10-63 | 80 | A |
| 7710-M32-4-5-10-64 | 100 | D |
| 7710-M32-4-5-10-65 | 0 | A |
| 7710-M32-4-5-10-66 | 100 | D |
| 7710-M32-4-5-10-67 | 0 | A |
| 7710-M32-4-5-10-68 | 100 | D |
| 7710-M32-4-5-10-69 | 0 | A |
| 7710-M32-4-5-10-70 | 0 | A |
| 7710-M32-4-5-10-71 | 0 | A |
| 7710-M32-4-5-10-72 | 0 | A |
| 7710-M32-4-5-10-73 | 0 | A |
| 7710-M32-4-5-10-74 | 100 | D |
| 7710-M32-4-5-10-75 | 100 | D |
| 7710-M32-4-5-10-76 | 0 | A |
| 7710-M32-4-5-10-77 | 0 | A |
| 7710-M32-4-5-10-78 | 100 | D |
| 7710-M32-4-5-10-79 | 0 | A |

| | | |
|---|---|---|
| 7710-M32-4-5-10-80 | 0 | A |
| 7710-M32-4-5-10-81 | 0 | A |
| 7710-M32-4-5-10-82 | 0 | A |
| 7710-M32-4-5-10-83 | 0 | A |
| 7710-M32-4-5-10-84 | 0 | A |
| 7710-M32-4-5-10-85 | 100 | D |
| 7710-M32-4-5-10-86 | 20 | A |
| 7710-M32-4-5-10-87 | 0 | A |
| 7710-M32-4-5-10-88 | 100 | D |
| 7710-M32-4-5-10-89 | 100 | D |
| 7710-M32-4-5-10-90 | 100 | D |
| 7710-M32-4-5-10-91 | | No data |
| 7710-M32-4-5-10-92 | 0 | A |
| 7710-M32-4-5-10-93 | 0 | A |
| 7710-M32-4-5-10-94 | 30 | A |
| 7710-M32-4-5-10-95 | 100 | D |
| 7710-M32-4-5-10-96 | 100 | D |
| 7710-M32-4-5-10-97 | 20 | A |
| 7710-M32-4-5-10-98 | 20 | A |
| 7710-M32-4-5-10-99 | 100 | D |
| 7710-M32-4-5-10-100 | 20 | A |
| 7710-M32-4-5-10-101 | 20 | A |
| 7710-M32-4-5-10-102 | 20 | A |
| 7710-M32-4-5-10-103 | 20 | A |
| 7710-M32-4-5-10-104 | 20 | A |
| 7710-M32-4-5-10-105 | 20 | A |
| 7710-M32-4-5-10-106 | 20 | A |
| 7710-M32-4-5-10-107 | 100 | D |
| 7710-M32-4-5-10-108 | 20 | A |
| 7710-M32-4-5-10-109 | 80 | A |
| 7710-M32-4-5-10-110 | 20 | A |
| 7710-M32-4-5-10-111 | 20 | A |
| 7710-M32-4-5-10-112 | 20 | A |
| 7710-M32-4-5-10-113 | 20 | A |
| 7710-M32-4-5-10-114 | 80 | A |
| 7710-M32-4-5-10-115 | 100 | D |
| 7710-M32-4-5-10-116 | 20 | A |
| 7710-M32-4-5-10-117 | 20 | A |
| 7710-M32-4-5-10-118 | 20 | A |

| | | |
|---|---|---|
| 7710-M32-4-5-10-119 | 20 | A |
| 7710-M32-4-5-10-120 | 20 | A |
| 7710-M32-4-5-10-121 | 100 | D |
| 7710-M32-4-5-10-122 | 20 | A |
| 7710-M32-4-5-10-123 | 100 | D |
| 7710-M32-4-5-10-124 | 20 | A |
| 7710-M32-4-5-10-125 | 20 | A |
| 7710-M32-4-5-10-126 | 20 | A |
| 7710-M32-4-5-10-127 | 100 | D |
| 7710-M32-4-5-10-128 | 100 | D |
| 7710-M32-4-5-10-129 | 100 | D |
| 7710-M32-4-5-10-130 | 0 | A |
| 7710-M32-4-5-10-131 | 0 | A |
| 7710-M32-4-5-10-132 | 80 | A |
| 7710-M32-4-5-10-133 | 20 | A |
| 7710-M32-4-5-10-134 | | No data |
| 7710-M32-4-5-10-135 | 0 | A |
| 7710-M32-4-5-10-136 | 0 | A |
| 7710-M32-4-5-10-137 | 100 | D |
| 7710-M32-4-5-10-138 | | No data |
| 7710-M32-4-5-10-139 | 100 | D |
| 7710-M32-4-5-10-140 | 100 | D |
| 7710-M32-4-5-10-141 | | No data |
| 7710-M32-4-5-10-142 | | No data |
| 7710-M32-4-5-10-143 | | No data |
| 7710-M32-4-5-10-144 | | No data |
| 7710-M32-4-5-10-145 | | No data |
| 7710-M32-4-5-10-146 | 100 | D |
| 7710-M32-4-5-10-147 | | No data |
| 7710-M32-4-5-10-148 | 80 | A |
| 7710-M32-4-5-10-149 | | No data |
| 7710-M32-4-5-10-150 | 100 | D |
| 7710-M32-4-5-10-151 | | No data |
| 7710-M32-4-5-10-152 | 80 | A |
| 7710-M32-4-5-10-153 | | No data |
| 7710-M32-4-5-10-154 | | No data |
| 7710-M32-4-5-10-155 | 80 | A |
| 7710-M32-4-5-10-156 | | No data |
| 7710-M32-4-5-10-157 | 100 | D |

| | | |
|---|---|---|
| 7710-M32-4-5-10-158 | 80 | A |
| 7710-M32-4-5-10-159 | 100 | D |
| 7710-M32-4-5-10-160 | 30 | A |
| 7710-M32-4-5-10-161 | 100 | D |
| 7710-M32-4-5-10-162 | 70 | A |
| 7710-M32-4-5-10-163 | 100 | D |
| 7710-M32-4-5-10-164 | 100 | D |
| 7710-M32-4-5-10-165 | 70 | A |
| 7710-M32-4-5-10-166 | 70 | A |
| 7710-M32-4-5-10-167 | 100 | D |
| 7710-M32-4-5-10-168 | 100 | D |
| 7710-M32-4-5-10-169 | 70 | A |
| 7710-M32-4-5-10-170 | 100 | D |
| 7710-M32-4-5-10-171 | | No data |
| 7710-M32-4-5-10-172 | 70 | A |
| 7710-M32-4-5-10-173 | 30 | A |
| 7710-M32-4-5-10-174 | 70 | A |
| 7710-M32-4-5-10-175 | 30 | A |
| 7710-M32-4-5-10-176 | 30 | A |
| 7710-M32-4-5-10-177 | 30 | A |
| 7710-M32-4-5-10-178 | | No data |
| 7710-M32-4-5-10-179 | 100 | D |
| 7710-M32-4-5-10-180 | 30 | A |
| 7710-M32-4-5-10-181 | 100 | D |
| 7710-M32-4-5-10-182 | 100 | D |
| 7710-M32-4-5-10-183 | 30 | A |
| 7710-M32-4-5-10-184 | 100 | D |
| 7710-M32-4-5-10-185 | 30 | A |
| 7710-M32-4-5-10-186 | | No data |
| 7710-M32-4-5-10-187 | | No data |
| 7710-M32-4-5-10-188 | 30 | A |
| 7710-M32-4-5-10-189 | 100 | D |
| 7710-M32-4-5-10-190 | 30 | A |
| 7710-M32-4-5-10-191 | 20 | A |
| 7710-M32-4-5-10-192 | 70 | A |
| 7710-M32-4-5-10-193 | 100 | D |
| 7710-M32-4-5-10-194 | 80 | A |
| 7710-M32-4-5-10-195 | | No data |
| 7710-M32-4-5-10-196 | | No data |

| | | |
|---|---|---|
| 7710-M32-4-5-10-197 | 30 | A |
| 7710-M32-4-5-10-198 | | No data |
| 7710-M32-4-5-10-199 | | No data |
| 7710-M32-4-5-10-200 | | No data |
| 7710-M32-4-5-10-201 | 30 | A |
| 7710-M32-4-5-10-202 | 80 | A |
| 7710-M32-4-5-10-203 | | No data |
| 7710-M32-4-5-10-204 | 100 | D |
| 7710-M32-4-5-10-205 | | No data |
| 7710-M32-4-5-10-206 | 100 | D |
| 7710-M32-4-5-10-207 | 30 | A |
| 7710-M32-4-5-10-208 | 100 | D |
| 7710-M32-4-5-10-209 | | No data |
| 7710-M32-4-5-10-210 | 30 | A |
| 7710-M32-4-5-10-211 | 30 | A |
| 7710-M32-4-5-10-212 | 30 | A |
| 7710-M32-4-5-10-213 | 100 | D |
| 7710-M32-4-5-10-214 | 30 | A |
| 7710-M32-4-5-10-215 | 100 | D |
| 7710-M32-4-5-10-216 | 30 | A |
| 7710-M32-4-5-10-217 | 30 | A |
| 7710-M32-4-5-10-218 | 30 | A |
| 7710-M32-4-5-10-219 | 30 | A |
| 7710-M32-4-5-10-220 | 100 | D |
| 7710-M32-4-5-10-221 | 100 | D |
| 7710-M32-4-5-10-222 | 30 | A |
| 7710-M32-4-5-10-223 | 100 | D |
| 7710-M32-4-5-10-224 | 80 | A |
| 7710-M32-4-5-10-225 | 20 | A |
| 7710-M32-4-5-10-226 | 100 | D |
| 7710-M32-4-5-10-227 | 100 | D |
| 7710-M32-4-5-10-228 | | No data |
| 7710-M32-4-5-10-229 | 20 | A |
| 7710-M32-4-5-10-230 | | No data |
| 7710-M32-4-5-10-231 | 20 | A |
| 7710-M32-4-5-10-232 | 100 | D |
| 7710-M32-4-5-10-233 | | No data |
| 7710-M32-4-5-10-234 | 0 | A |
| 7710-M32-4-5-10-235 | | No data |

| | | |
|---|---|---|
| 7710-M32-4-5-10-236 | | No data |
| 7710-M32-4-5-10-237 | 100 | D |
| 7710-M32-4-5-10-238 | 0 | A |
| 7710-M32-4-5-10-239 | 0 | A |
| 7710-M32-4-5-10-240 | 80 | A |
| 7710-M32-4-5-10-241 | 30 | A |
| 7710-M32-4-5-10-242 | | No data |
| 7710-M32-4-5-10-243 | 0 | A |
| 7710-M32-4-5-10-244 | 100 | D |
| 7710-M32-4-5-10-245 | | No data |
| 7710-M32-4-5-10-246 | | No data |
| 7710-M32-4-5-10-247 | 0 | A |
| 7710-M32-4-5-10-248 | 100 | D |
| 7710-M32-4-5-10-249 | 100 | D |
| 7710-M32-4-5-10-250 | 0 | A |
| 7710-M32-4-5-10-251 | 30 | A |
| 7710-M32-4-5-10-252 | 0 | A |
| 7710-M32-4-5-10-253 | 0 | A |
| 7710-M32-4-5-10-254 | 0 | A |
| 7710-M32-4-5-10-255 | 100 | D |
| 7710-M32-4-5-10-256 | 0 | A |
| 7710-M32-4-5-10-257 | 0 | A |
| 7710-M32-4-5-10-258 | 100 | D |
| 7710-M32-4-5-10-259 | 100 | D |
| 7710-M32-4-5-10-260 | 100 | D |
| 7710-M32-4-5-10-261 | 100 | D |
| 7710-M32-4-5-10-262 | 100 | D |
| 7710-M32-4-5-10-263 | 0 | A |
| 7710-M32-4-5-10-264 | 100 | D |
| 7710-M32-4-5-10-265 | 80 | A |
| 7710-M32-4-5-10-266 | 80 | A |
| 7710-M32-4-5-10-267 | 0 | A |
| 7710-M32-4-5-10-268 | | No data |
| 7710-M32-4-5-10-269 | 100 | D |
| 7710-M32-4-5-10-270 | 0 | A |
| 7710-M32-4-5-10-271 | 100 | D |
| 7710-M32-4-5-10-272 | 20 | A |
| 7710-M32-4-5-10-273 | 0 | A |
| 7710-M32-4-5-10-274 | 80 | A |

| | | |
|---|---|---|
| 7710-M32-4-5-10-275 | 100 | D |
| 7710-M32-4-5-10-276 | 100 | D |
| 7710-M32-4-5-10-277 | 30 | A |
| 7710-M32-4-5-10-278 | 100 | D |
| 7710-M32-4-5-10-279 | 80 | A |
| 7710-M32-4-5-10-280 | 100 | D |
| 7710-M32-4-5-10-281 | 100 | D |
| 7710-M32-4-5-10-282 | 100 | D |
| 7710-M32-4-5-10-283 | 100 | D |
| 7710-M32-4-5-10-284 | 20 | A |
| 7710-M32-4-5-10-285 | 0 | A |
| 7710-M32-4-5-10-286 | 0 | A |
| 7710-M32-4-5-10-287 | | No data |
| 7710-M32-4-5-10-288 | 100 | D |
| 7710-M32-4-5-10-289 | 80 | A |
| 7710-M32-4-5-10-290 | | No data |
| 7710-M32-4-5-10-291 | 100 | D |
| 7710-M32-4-5-10-292 | 30 | A |
| 7710-M32-4-5-10-293 | 30 | A |
| 7710-M32-4-5-10-294 | 30 | A |
| 7710-M32-4-5-10-295 | 100 | D |
| 7710-M32-4-5-10-296 | 100 | D |
| 7710-M32-4-5-10-297 | 30 | A |
| 7710-M32-4-5-10-298 | 30 | A |
| 7710-M32-4-5-10-299 | 30 | A |
| 7710-M32-4-5-10-300 | 30 | A |
| 7710-M32-4-5-10-301 | 30 | A |
| 7710-M32-4-5-10-302 | 30 | A |
| 7710-M32-4-5-10-303 | 30 | A |
| 7710-M32-4-5-10-304 | 100 | D |
| 7710-M32-4-5-10-305 | 100 | D |
| 7710-M32-4-5-10-306 | 30 | A |
| 7710-M32-4-5-10-307 | 30 | A |
| 7710-M32-4-5-10-308 | 100 | D |
| 7710-M32-4-5-10-309 | 100 | D |
| 7710-M32-4-5-10-310 | 30 | A |
| 7710-M32-4-5-10-311 | 30 | A |
| 7710-M32-4-5-10-312 | 30 | A |
| 7710-M32-4-5-10-313 | 30 | A |

| | | |
|---|---|---|
| 7710-M32-4-5-10-314 | | No data |
| 7710-M32-4-5-10-315 | 30 | A |
| 7710-M32-4-5-10-316 | 30 | A |
| 7710-M32-4-5-10-317 | 100 | D |
| 7710-M32-4-5-10-318 | 100 | D |
| 7710-M32-4-5-10-319 | 30 | A |
| 7710-M32-4-5-10-320 | 30 | A |
| 7710-M32-4-5-10-321 | 100 | D |
| 7710-M32-4-5-10-322 | 80 | A |
| 7710-M32-4-5-10-323 | 100 | D |
| 7710-M32-4-5-10-324 | | No data |
| 7710-M32-4-5-10-325 | 0 | A |
| 7710-M32-4-5-10-326 | 100 | D |
| 7710-M32-4-5-10-327 | 100 | D |
| 7710-M32-4-5-10-328 | 100 | D |
| 7710-M32-4-5-10-329 | 100 | D |
| 7710-M32-4-5-10-330 | 20 | A |
| 7710-M32-4-5-10-331 | | No data |
| 7710-M32-4-5-10-332 | 80 | A |
| 7710-M32-4-5-10-333 | 80 | A |
| 7710-M32-4-5-10-334 | 0 | A |
| 7710-M32-4-5-10-335 | 0 | A |
| 7710-M32-4-5-10-336 | 30 | A |
| 7710-M32-4-5-10-337 | 30 | A |
| 7710-M32-4-5-10-338 | 100 | D |
| 7710-M32-4-5-10-339 | 100 | D |
| 7710-M32-4-5-10-340 | 0 | A |
| 7710-M32-4-5-10-341 | 0 | A |
| 7710-M32-4-5-10-342 | 0 | A |
| 7710-M32-4-5-10-343 | 0 | A |
| 7710-M32-4-5-10-344 | 0 | A |
| 7710-M32-4-5-10-345 | 0 | A |
| 7710-M32-4-5-10-346 | 0 | A |
| 7710-M32-4-5-10-347 | 100 | D |
| 7710-M32-4-5-10-348 | 0 | A |
| 7710-M32-4-5-10-349 | 80 | A |
| 7710-M32-4-5-10-350 | 20 | A |
| 7710-M32-4-5-10-351 | 0 | A |
| 7710-M32-4-5-10-352 | 80 | A |
